# Universal bacterial clade dynamics dominate under predation despite altered phenotypes and mutation targets

**DOI:** 10.1101/2025.02.04.636436

**Authors:** Dovydas Kičiatovas, Johannes Cairns, Veera Partanen, Julius Hoffmann, Lutz Becks, Teppo Hiltunen, Ville Mustonen

**Author notes:** Corresponding authors: Lutz Becks,; Teppo Hiltunen,; Ville Mustonen.

## Abstract

Recent studies have revealed bacterial genome-wide evolution to be complex and dynamic even in a constant environment, characterized by the emergence of new clades competing or temporarily coexisting as each clade undergoes evolutionary change. Previous studies on predator-prey dynamics tracking simple ecological and phenotypic metrics have shown predation to fundamentally alter prey evolution, facilitating defense evolution followed by coevolution and frequency dependent selection between defended and undefended prey genotypes. Here we sought to consolidate these fields by examining genome-wide evolution in five bacterial prey species separately subjected to long-term evolution under ciliate predation. We hypothesized that the presence of predation could change the pattern of clonal dynamics, for example, by more frequently producing selective sweeps if predation-defense-related mutations are under strong selection. For all species, we found mutational signals of prey adaptation, with phenotypic data and genomic mutation targets demonstrating changes in composition between the experimental treatments. Intriguingly, despite higher variant counts, overall temporal clade dynamics across the coevolved prey species were strikingly similar to those of bacteria evolving alone, with constant emergence, competition and quasi-stable coexistence of clades. This study shows that long-term molecular evolution in bacterial prey under predation is more interesting and less predictable than we might expect based on existing coevolutionary theories.

## Introduction

Understanding how organisms adapt and evolve is a fundamental challenge in biology that continues to intrigue researchers. While organisms seemingly respond to environmental pressures through predictable phenotypic changes (Stern 2013), the underlying genetic mechanisms reveal a far more complex and unpredictable reality (Tenaillon et al. 2012). At the phenotypic level, adaptation is often repeatable and predictable, both in controlled experiments (Wichman et al. 1999; Katz et al. 2021) and when comparing across natural populations (Papadopulos et al. 2021). This repeatability can allow us to identify key mechanisms driving adaptive changes, such as selection strength (Saxer, Doebeli, and Travisano 2010; Bernardes et al. 2021) and trade-offs between traits (Soudi et al. 2023; Maccagni and Willi 2022). However, at the nucleotide level, evolution is typically much less repeatable, even under strong selective pressures (Lässig, Mustonen, and Walczak 2017). This lack of repeatability arises from two main factors. First, different metabolic pathways can lead to the same phenotypic outcome, allowing for multiple routes of adaptation (Frickel et al. 2018; Maharjan et al. 2015). Second, even in relatively simple environments and within a single, isolated population, clonal structure can undergo continuous and significant changes over time. These changes are driven by complex dynamics, such as competition and persistence among mutational or clonal cohorts, which can coexist for extended periods (Good et al. 2017; Lang et al. 2013). Clonal cohorts are groups of genetically identical individuals that arise from a common ancestor and share the same mutations. These cohorts play a crucial role in shaping the genetic composition of populations over time, as they compete, persist, or decline based on their fitness advantages or disadvantages, and chance. Thus, molecular evolution is shaped not only by direct selection and the emergence of new mutations in individuals constituting the populations, but also by non-trivial interactions among clonal cohorts within populations over generations. Understanding these dynamics is crucial for uncovering how genetic changes respond to environmental pressures and why evolutionary outcomes can diverge, even under similar selective conditions.

Ecological interactions, those between individuals of different species, introduce additional layers of complexity to molecular evolution (Cairns et al. 2020b). For instance, species interactions can drive strong directional or stabilizing selection on populations. Strong selection may increase the efficiency of eliminating competing clonal lineages, simplifying clonal population dynamics (Corbett-Detig, Hartl, and Sackton 2015). On the other hand, species interactions can lead to coevolution and eco-evolutionary dynamics, where changes in population size both drive and result from evolutionary shifts occurring on similar timescales (Lion 2018). In these scenarios, selection pressures fluctuate continuously, varying in direction and/or strength. These fluctuations can promote the coexistence of clonal cohorts in a quasi-stable or frequency-dependent manner, making molecular evolution at least as dynamic as in single-species systems. For example, a study of microbial host-virus coevolution found that the host harbored significant genetic diversity, primarily driven by neutral molecular evolution rather than direct selection by the virus. The virus underwent fewer, strongly selected mutations, typically confined to specific genes (Retel et al. 2019). Furthermore, the complexity of species interactions can lead to conflicting selection pressures, potentially altering the trajectory of molecular evolution. This has been observed in tripartite interactions involving bacteria, phages, and plasmids, where the presence of multiple mobile genetic elements constrains the evolutionary responses typically seen in pairwise interactions (Harrison et al. 2017). Coevolution can also accelerate molecular evolution (Paterson et al. 2010), driving successive sweeps of beneficial mutations and, in some cases, leading to simpler evolutionary dynamics. These examples illustrate how coevolutionary dynamics can lead to both complex or more simple patterns of molecular evolution.

Predator-prey relationships are commonly encountered in nature and represent important examples of eco-evolutionary dynamics. Although predation overall is an important selective pressure (Yoshida et al. 2003), the strength of selection and breadth of genomic targets are likely to substantially vary depending on the predator-prey system, i.e., generalist vs. specialist predator and parameters such as predator feeding rate (Filip et al. 2014), and the genomic and physiological constraints for anti-predatory defense mechanisms (Matz and Kjelleberg 2005). Furthermore, adaptation to the growth environment and available resources is critical for both competitive and defensive capabilities of prey microbes (Friman et al. 2008), meaning that the prey evolution might be subject to both strong biotic (predator) and strong abiotic selective pressures. Therefore, we chose to study a known predator-prey system – ciliate (*Tetrahymena thermophila*) preying upon bacterial prey species – in which strong phenotypic and genomic changes have been previously observed as the outcome of in vitro coevolution, making it an ideal system to test for detectable molecular evolutionary signals (Cairns et al. 2020a). *T. thermophila* grazes upon bacterial populations by ingesting cells through the oral apparatus and mainly selects prey based on physical dimensions (Hahn and Höfle 2001). Thus, common ciliate prey defense mechanisms include motility, oversize and biofilm formation (Matz and Kjelleberg 2005). Previous studies on this ciliate-bacteria system have shown that ciliate predation is a strong selective pressure, inducing rapid adaptation in prey (Hiltunen and Becks 2014; Friman, Jousset, and Buckling 2014).

Furthermore, the patterns and rates of molecular evolution differ depending on the timescale observed. Experimental evolution studies, which track molecular evolution in replicated populations, have shown that, over short timescales, beneficial mutations accumulate rapidly in response to new selective pressures. These mutations are often large-effect and lead to significant fitness gains in a short period, allowing for an evaluation of predictability in evolutionary processes (Johnson et al. 2021). Such acceleration in genome evolution was shown in other predator-prey systems, such as in myxobacterium preying on *E. coli* (Nair et al. 2019). Long-term dynamics, however, are more likely governed by declining rates of adaptation, as beneficial mutations become rarer, while the overall rate of molecular evolution remains constant. Although much of our understanding of molecular evolution in the context of species interactions comes from short-term experiments, observation over longer periods exhibits evolutionary dynamics that are not visible over short time scales (Good et al. 2017). Unfortunately, such long-term studies are currently lacking in terms of species interactions, leaving unanswered questions about how they influence molecular evolution. Understanding these dynamics is essential for predicting and managing evolutionary outcomes in natural and artificial systems, from ecosystem conservation to the development of sustainable disease control strategies (Lässig, Mustonen, and Nourmohammad 2023). In this study, we sought in particular to investigate whether antagonistic predator-prey relationships leave a lasting molecular signature in bacterial prey evolving under constant predation.

To that end, we investigated the long-term molecular and phenotypic evolution of five bacterial species (*Brevundimonas diminuta*, *Comamonas testosteroni*, *Pseudomonas fluorescens*, *Sphingomonas capsulata*, and *Serratia marcescens*) as they grew with and without a ciliate predator, *Tetrahymena thermophila*, over a period of 800 days (Fig. 1, first two panels; see Methods). Our study builds upon the long-term experimental evolution methodology presented in Good et al. 2017, where bacterial molecular evolution is shown to be complex and incessant over long time scales, even in a constant environment. The five focal prey species in this study are relevant to predator-prey studies due to various characteristics: *B. diminuta* has been previously studied and is found in lake habitats shared with Tetrahymena (Becks et al. 2005), *C. testosteroni* is known to have specific defenses against protozoan grazing (Matz and Kjelleberg 2005). The remaining three – *P. fluorescens*, *S. capsulata* and *S. marcescens* – are model prey species previously used in predator–prey studies (Hiltunen and Laakso 2013; Hiltunen et al. 2018). For each species and treatment combination, three replicate populations were evolved – resulting in 30 populations. This long-term experiment is a direct continuation of the predator-prey evolutionary experiment described in Cairns et al. 2020a – in this study, we focused on a subset of five species instead of seven described in the original study because of experimental issues related to the remaining two species (see Methods). Here, along with similar phenotypic data collection like in the original study, we collected and sequenced whole genomes of each evolving population, at intervals of 28 days (see Methods). After accounting for unused samples (due to contamination, low quality, etc.), each study population is represented by up to 25 time-resolved genomic samples, spanning 800 days of evolution. Our aim was to quantitatively assess how the presence of the predator influenced molecular evolution in the bacterial prey (Fig. 1, third panel; see Methods). We sought to determine whether the predator’s presence is detectable in the clonal structure of prey populations and growth characteristics, during and after long-term evolution, by comparing prey populations coevolving with the ciliate versus populations evolving alone. Since we know from earlier studies of this predator-prey system that ciliate presence affects growth phenotypes and causes non-random recurrent mutations (Cairns et al. 2020a), we hypothesized that its presence would also be reflected in the clonal dynamics of the prey – e.g., by increasing the occurrence of selective sweeps if certain mutations targeting crucial defense genes are under strong selection. To investigate this, we tracked clonal diversity and phenotypic changes in replicated bacterial populations over time, and identified intraspecific recurrence of mutation gene targets.

**Figure 1:**
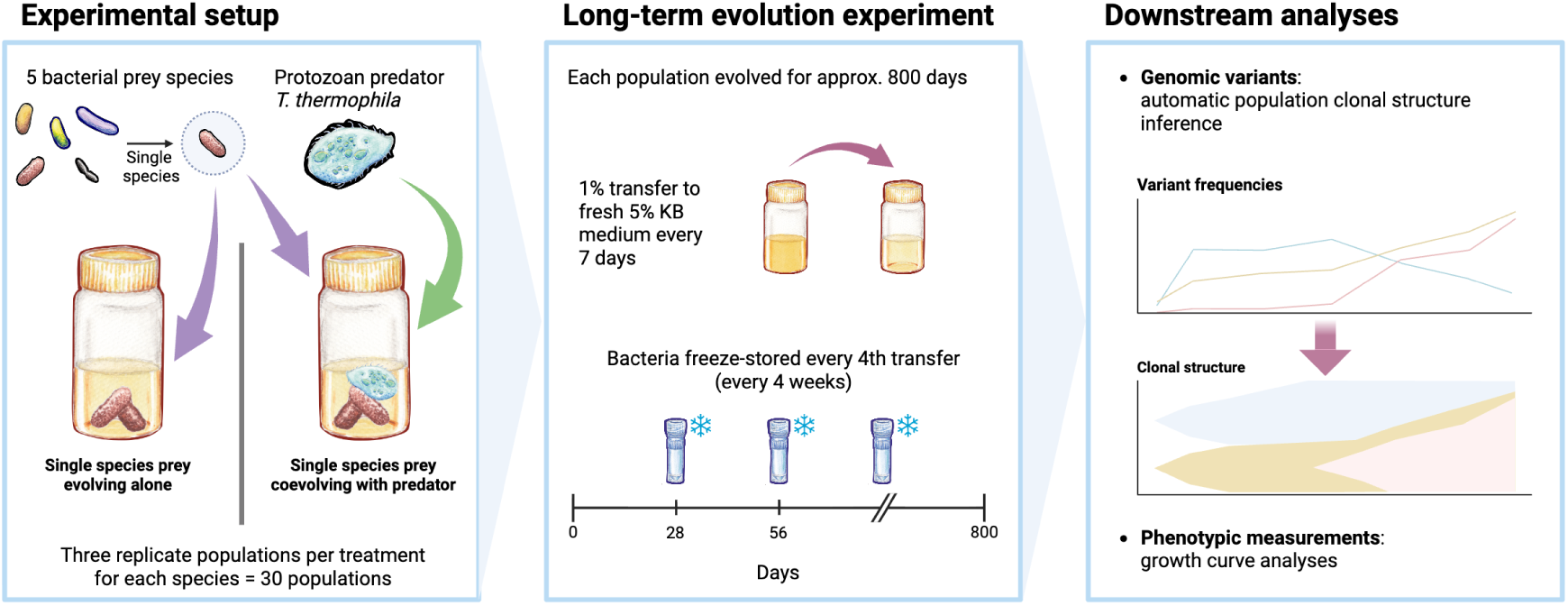
Graphical abstract of this study. Each of the five prey species were evolved along two tracks of long-term evolution – evolved alone and coevolved with predator *T. thermophila*. Each prey population was freeze-stored every four transfers (i.e., weeks), and later sampled to extract bacterial DNA for genomic sequencing (see Methods). Additionally, isolates for phenotypic measurements were taken twice in the experiment (early and late; see Methods). Therefore, the resulting genomic and phenotypic data are time-resolved and allow for clonal structure inferences.

## Results

### Growth phenotypes differ between prey populations coevolving with a predator and populations evolving alone

First, to verify that the ciliate has exerted sufficient evolutionary pressure to induce evolutionary adaptation in the bacterial prey during the long-term experiment, we analysed the growth curve data (i.e., growth phenotypes) from samples of each study population at the beginning and the end of the long-term experiment, as well as from ancestral samples (see Methods). Prey that has evolved alone should exhibit lower growth when exposed to the ciliate than prey that has coevolved with the ciliate. The same pattern should be detectable when comparing coevolved isolates with their ancestors. To that end, we compared growth characteristics between prey populations that evolved alone and populations that coevolved with the ciliate (hereafter we collectively refer to both treatment lines as evolutionary histories), and also contrasted isolates of both evolutionary histories against their species-specific ancestor isolates.

Here, we sought to describe the phenotypic changes in these prey species over a longer period of time than previously studied. Adaptation to a particular stressor typically influences multiple bacterial phenotypes due to the pleiotropic effects of altering key structures or functions, as is the case with antibiotic cross resistance and collateral sensitivity (Colclough et al. 2019). Therefore, in addition to studying the phenotypes of bacteria evolved alone or with ciliate in the primary conditions of interest – experimental medium with and without ciliate predation – we also included a salt stress condition to further measure a fitness cost of prey of either treatment lineage when grown without the predator. Thus, the sampled prey population isolates were grown on three nutrient-rich experimental media – without ciliate (control), with ciliate and with added KCl salt without ciliate (see Methods).

For each growth curve, we computed yield *Y* (final population size), the area under the curve (AUC; the integral of the entire growth curve, i.e., total number of cells produced over the duration of growth), maximal per-capita growth rate *ρ_max_*, and time to the latter *t_ρ__max_* (see Methods). For subsequent analyses, we chose AUC as the representative growth curve statistic, as it is commonly used to summarize the entire growth curve in one value, allowing for easy processing and comparisons. The remaining three metrics – *Y*, *ρ_max_*, and *t_ρ__max_* – may be more susceptible to the measurement noise and they use only a part of the curve. We did not observe a consistent direction of growth phenotype evolution with respect to time in the long-term experiment. Isolates sampled later in the experiment may grow better or worse in terms of AUC than in the beginning without a clear pattern across replicates, though it clearly differs between species (Supplementary Fig. 1). Despite the overall signal separating evolutionary histories in terms of differences in AUC in the presence of predation, each species has a unique phenotypic response to each growth medium, regardless of their evolutionary history. Supplementary Figs. 2, 3 and 4 show similar box plots of isolate yield *Y*, *ρ_max_*and *t_ρ__max_*, respectively.

To determine whether predation exerted a sufficiently strong selective pressure to affect prey population growth dynamics in our experiment, we computed ratios of AUC between species-specific ancestral and (co-)evolved population growth curves (see Methods; Fig. 2). Then, we fitted a linear mixed model to data of each of the three growth media separately to investigate the growth differences based on species identity, evolutionary history and sampling timing (the main fixed terms), and pairwise interactions between them; to account for repeated measurements of the same replicates over time, we modeled replicates as random effects (see Methods). Additionally, we quantified the contribution of the main fixed terms to the observed phenotypic patterns in terms of proportion of variance explained (Fig. 2, right hand panels; see Methods). We then tested whether the addition of various interactions between our base linear mixed model terms results in improved models. Using a log-likelihood ratio test, we tested the inclusion of three interactions: between evolutionary history and species identity, between evolutionary history and sampling timing, and between species identity and sampling timing (see Methods). Including all three interactions as fixed effects results confers a significantly better model fit (Supplementary Fig. 5; model *R*^2^ and log-likelihood ratio test *χ*^2^(extra degrees of freedom) and *p* values: control medium base model *R*^2^ = 0.42, interaction model *R*^2^ = 0.62, ratio test *χ*^2^(9) = 255.66, *p <* 0.001; salt stress medium base model *R*^2^ = 0.36, interaction model *R*^2^ = 0.54, ratio test *χ*^2^(9) = 210.5, *p <* 0.001; predator medium base model *R*^2^ = 0.61, interaction model *R*^2^ = 0.71, ratio test *χ*^2^(9) = 190.83, *p <* 0.001).

**Figure 2:**
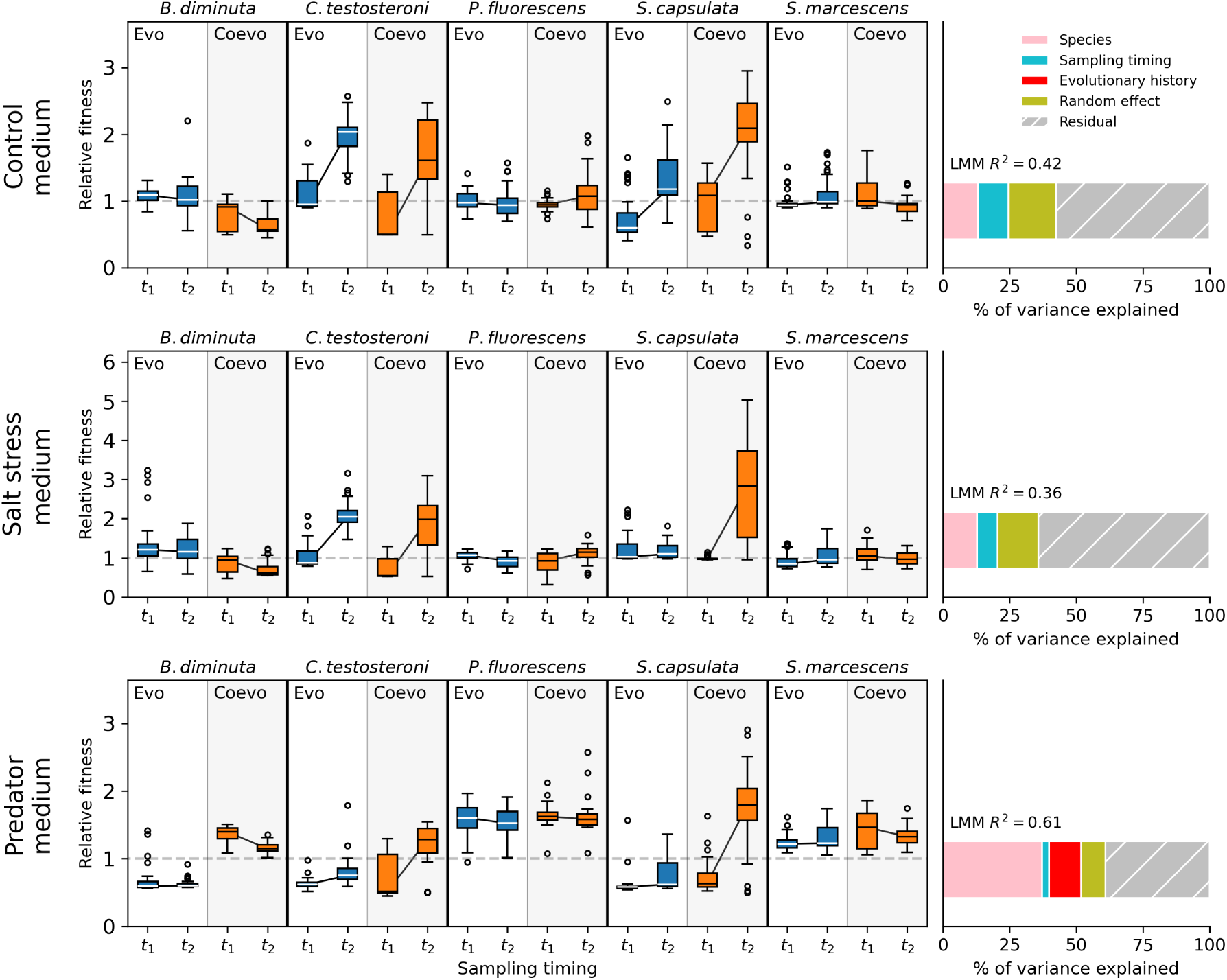
Fitness relative to the ancestor in terms of area under the curve (AUC) ratios between evolved isolates from populations of each study species and ancestral isolates of the respective species (see Methods). Each panel (row) represents one of the three growth media and is divided into five sections: one for each study species (see Methods). Each box plot represents the phenotypic sampling timing (*t*_1_: early, *t*_2_: late) in the three populations of each species (each box represented by 30 isolates), which are denoted *Evo* for evolved-alone populations (represented with blue boxes) and *CoEvo* for populations that coevolved with the ciliate (represented with orange boxes). For boxes representing evolutionary histories within each species, a black line connects the data medians of its two sampling time points. A dashed horizontal line represents no change in fitness (AUC ratio = 1). A panel on the right hand side of each row shows the proportion of variance explained by each of the main terms (species identity, evolutionary history, sampling timing) with the random term in the base (i.e., without interactions between main terms) linear mixed model fitted on fitness data of each medium. Hatched gray area represents the unexplained (residual) variance. Total proportion of explained variance of the base linear mixed model, represented by *R*^2^ value, is shown above each bar.

We saw a varied response to predation across the species, both in strength and direction with respect to isolate sampling timing (Fig. 2, black lines between early and late isolate boxes). The bacterium were expected to evolve anti-predatory defenses in the presence of the ciliate, as shown before in a previously described ciliate-*P. fluorescens* system in Scheuerl *et al*. study (Scheuerl et al. 2019). However, interestingly, our *P. fluorescens* and also *S. marcescens* appear to have not been greatly affected by the predator, as even the evolved-alone populations have grown in the presence of the predator similarly to coevolved populations (Fig. 2; *P. fluorescens* and *S. marcescens* in “Predator medium” sub-panel). Nevertheless, in the remaining species, we saw the expected differences in evolutionary outcomes when grown with the predator present (predator medium data, log-likelihood ratio test *χ*^2^(extra degrees of freedom) and *p* values; evolutionary history: *χ*^2^(1) = 25.641, *p <* 0.001, species identity: *χ*^2^(4) = 49.354, *p <* 0.001, sampling timing: *χ*^2^(1) = 37.493, *p <* 0.001, evolutionary history and species identity interaction: *χ*^2^(4) = 33.345, *p <* 0.001, evolutionary history and sampling timing interaction: *χ*^2^(1) = 15.552, *p <* 0.001, species identity and sampling timing interaction: *χ*^2^(4) = 137.61, *p <* 0.001). The evolutionary history is a useful predictor only in the predator medium, contributing ∼12% of total explained variance, whereas it is *<*0.2% in the other media (Fig. 2, bottom row, right hand panel, red-shaded area). This means that the presence of ciliate resulted in the coevolved bacterium having a noticeably different phenotypic response compared to the bacterium that were evolved alone, and this difference is consistently observed in the medium where the ciliate is present. The growth phenotype differences between bacterial species were further exacerbated in the presence of the ciliate, as supported by the nearly three times more of variance explained by the species identity in the predator medium than in the other two media – ∼37% compared to ∼13% in both control and salt stress media.

It is noteworthy that most coevolved bacteria populations (a notable exception is *B. diminuta*) in general did not display lower AUC in the control medium, compared to the bacteria that evolved alone (control medium data, log-likelihood ratio test *χ*^2^(extra degrees of freedom) and *p* values; evolutionary history: *χ*^2^(1) = 0.2803, *p* = 0.5965, species identity: *χ*^2^(4) = 17.566, *p* = 0.0015, sampling timing: *χ*^2^(1) = 107.22, *p <* 0.001, evolutionary history and species identity interaction: *χ*^2^(4) = 21.522, *p <* 0.001, evolutionary history and sampling timing interaction: *χ*^2^(1) = 0.2676*, p* = 0.605, species identity and sampling timing interaction: *χ*^2^(4) = 233.73*, p <* 0.001). This could be explained by the expectation that coevolved prey populations would include both more reproductionally fit undefended phenotypes and ciliate-resistant but less fit defended phenotypes, as well as possibly both defended and more fit phenotypes, though that seems less likely due to the associated growth-defense trade-off (Huang et al. 2017). Furthermore, defensive trait expression could be triggered by the presence of a predator, resulting in a phenotypic change in cells that have defense-related genes but do not express them when predator is not present.

The coevolved *S. capsulata* populations seem to have gained a growth advantage in salt stress medium over the course of the experiment, as many of its isolates exhibited growth at the late sampling time point (*t*_2_), whereas the ancestors and early (*t*_1_) isolates did not grow in this medium (Fig. 2; *S. capsulata* “Salt stress” sub-panel; Supplementary Fig. 6). Otherwise, there seem to be no noticeable differences in growth ability between evolved-alone and coevolved prey (salt stress medium data, log-likelihood ratio test *χ*^2^(extra degrees of freedom) and *p* values; evolutionary history: *χ*^2^(1) = 0.0013, *p* = 0.9707, species identity: *χ*^2^(4) = 18.639, *p <* 0.001, sampling timing: *χ*^2^(1) = 67.888, *p <* 0.001, evolutionary history and species identity interaction: *χ*^2^(4) = 28.029, *p <* 0.001, evolutionary history and sampling timing interaction: *χ*^2^(1) = 22.233*, p <* 0.001, species identity and sampling timing interaction: *χ*^2^(4) = 153.2*, p <* 0.001).

Next, we used the random forest machine learning method to determine to what extent the study populations are distinguishable from one another in terms of their isolate growth. Each growth curve was collectively labeled with its corresponding species, evolutionary history, replicate population within the species and the timing of the isolate sampling. Then we concatenated the growth curves across the three types of media the isolates were grown in, and used that as input to train an ensemble of 100 random forest models with random splits of 50% of the data, predicted the label of the remaining randomly split 50% of growth curves and averaged the prediction accuracy across the random forest instances (see Methods; Fig. 3). As expected, bacterial adaptation in the presence or absence of ciliate predation had a multidimensional effect on bacterial phenotypes. Moreover, this effect was specific to the bacterial species, and within species, to the evolutionary history. However, the replicate populations are not as distinct within the pooled isolates of the same evolutionary history, and neither across the time points within a replicate population. Nevertheless, we see higher classification accuracy between the three replicate coevolved populations of each species (Fig. 3, right hand panel) than in populations that evolved without the predator, suggesting that the phenotypic dynamics are more distinguishable between coevolved populations.

**Figure 3:**
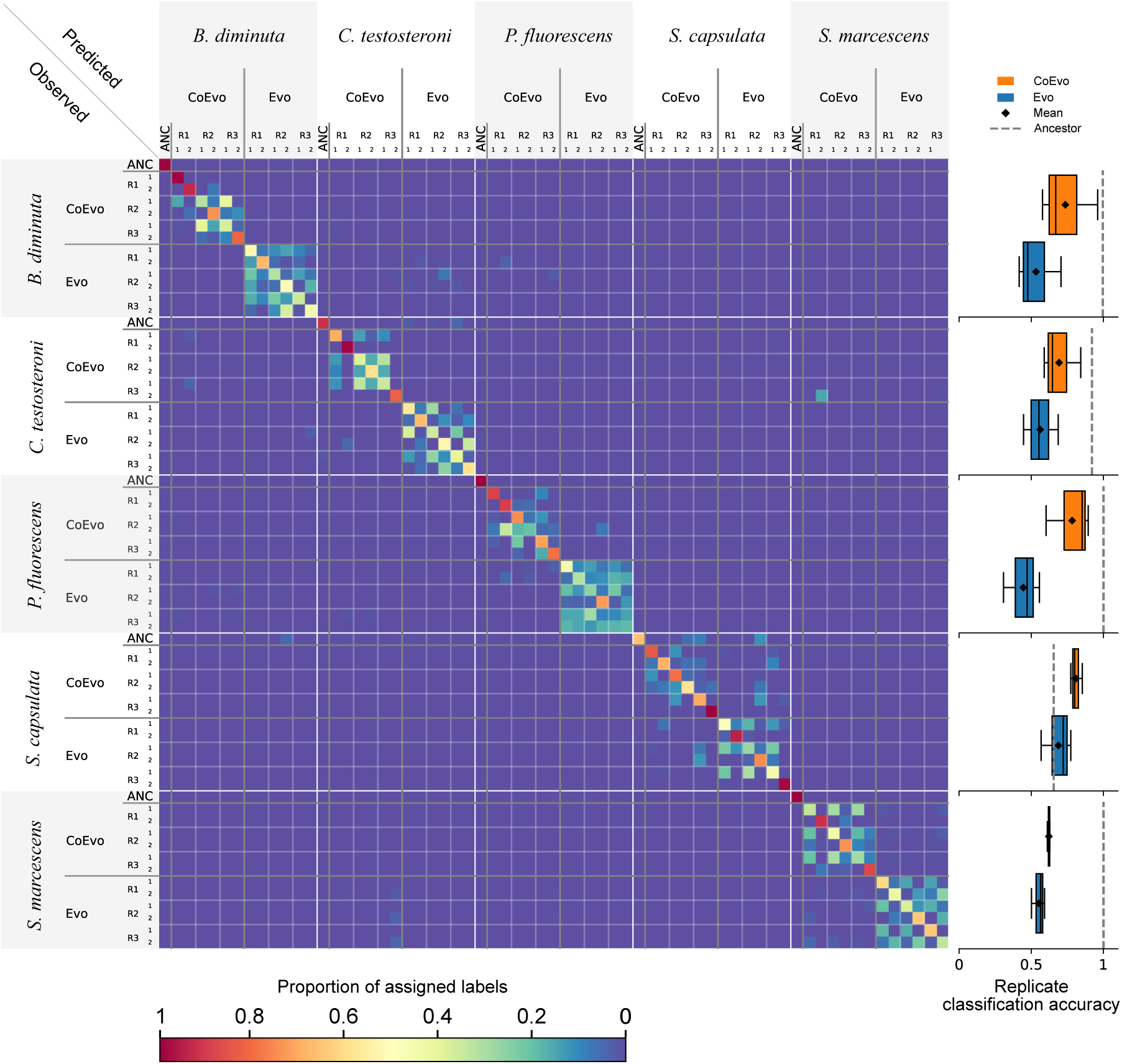
Combined and averaged confusion matrix of 100 random forest classifier instances, trained on 50% of randomly sampled isolate growth profiles (combined isolate growth curves in all three growth media) and tested on the remaining 50% of the data (label proportions preserved in both training and testing sets). The X-axis represents predicted data labels and the Y-axis – true labels. The rows and columns of the confusion matrix are separated according to the alphabetic order of the species names, and each inner square of the confusion matrix, representing a species, is further subdivided into the two evolutionary histories, then three replicate populations and lastly, the time points. “ANC” denotes the growth profile of the ancestral strain of the respective species. The color gradient represents the average number of assigned samples from the test set, averaged over 100 random forest instances. While there is little confusion between the species and evolutionary histories, the growth profiles of replicate populations within species and their evolutionary history are more difficult to separate. However, coevolved species-specific replicates are distinguished with higher accuracy than when evolved alone (right panel).

### Bacterial molecular evolutionary dynamics are similar regardless of the presence or absence of predator

Next, using the time-resolved sequencing data of each replicate of all prey species, evolving with and without the predator present, we calculated the frequencies of single nucleotide polymorphisms (SNPs) and short insertions or deletions (indels) and their change over the course of the experiment (see Methods). The final accumulated variant counts differ between species and between replicate populations (Supplementary Fig. 7, Supplementary Table 1), indicating a large degree of variability in the accumulated mutation counts even within the replicates of the same species under same environmental conditions. Likewise, population 1 of *C. testosteroni* steadily acquires almost double the number of variants compared to the remaining populations of either treatment of the same species. However, at the 95% confidence level, none of the species had significant differences when comparing the species-specific replicate means of final accumulated variant counts between evolutionary histories (two-sided Welch’s t-test; each group *n* = 3; *B. diminuta df* = 2.92, *p* = 0.94; *C. testosteroni df* = 2.83, *p* = 0.648; *P. fluorescens df* = 3.99, *p* = 0.545; *S. capsulata df* = 3.02, *p* = 0.696 and *S. marcescens df* = 2.00, *p* = 0.151). It is worth noting that the sample sizes for comparison are only three replicates per species per treatment – in future studies, larger sample sizes may allow for stronger inferences. The trajectories of most variants form clearly defined clusters, in line with the expectation that most mutations would hitchhike with beneficial mutations under selection (Lang and Desai 2014). Henceforth, we analyzed these clusters as variant cohorts (see Methods; Fig. 4A).

Several moderate to high-frequency variants in known mutator genes were observed together with a sharp increase in mutation counts right after the time of their detection, likely explaining why some study populations accrue much higher mutation counts than others within the same species and treatment. Variants in mutator gene *mutS* were detected in three species in both treatments: a stop-gain (appeared on day 700, mean frequency after appearance 0.32) and disruptive inframe deletion (appeared on day 476, mean frequency after appearance 0.36) in coevolved *P. fluorescens* populations 2 and 3, respectively; a missense variant in coevolved *S. marcescens* population 2 (appeared on day 679, mean frequency after appearance 0.75); disruptive (appeared on day 553, mean frequency after appearance 0.99) and conservative (appeared on day 553, mean frequency after appearance 0.96) inframe deletions in evolved-alone *C. testosteroni* populations 2 and 3, respectively; and a missense variant in evolved-alone *P. fluorescens* population 1 (appeared on day 412, mean frequency after appearance 0.64). Variants in *mutL* were detected in the following populations: a frameshift variant in coevolved *C. testosteroni* population 1 (appeared on day 448, mean frequency after appearance 0.35); a frameshift variant in evolved-alone *C. testosteroni* population 1 (appeared on day 146, mean frequency after appearance 0.66); and a stop-gain variant in evolved-alone *B. diminuta* population 1 (appeared on day 272, mean frequency after appearance 0.65). Additionally, in *C. testosteroni*, coevolved population 1 had a *uvrC* missense variant (appeared on day 504, mean frequency after appearance 0.13) and evolved-alone population 1 had two *uvrA* missense variants (appeared on days 553 and 776, mean frequencies after appearance 0.55 and 0.25).

Next, using the variant cohorts, we inferred the clonal structure within each population at every time point – treating the cohorts as genetic alleles, we developed a tree-based algorithm to assign each cohort to the role of either a new clone or a sub-clone of an existing clone (see Methods). Reconstructing clonal structures from variant frequency data is widely used in cancer research, where linking cancer evolution to clinical outcomes can inform therapies (Salcedo et al. 2020; Gillis and Roth 2020). Cancer clonal structure inference is difficult due to several factors, such as DNA copy number variation, genome diploidy and paucity of long term time-series data (Dentro et al. 2021), which are not present in our bacterial genomes, making this type of inference more readily viable. With the inferred allele hierarchy, we calculated the allele frequencies over time and visualize them as Muller plots and phylogenetic trees (Fig. 4A, B). Muller plots in this kind of population structure inference from variant data have been presented before (Frazão et al. 2022), however, to our knowledge, there is no tool that computes high-confidence population clonal structures automatically (nevertheless, we have manually checked all our visualised solutions as a form of quality control, see Methods).

We observed a large degree of variability in clonal dynamics, even between the replicates of the same species (Fig. 4, “Coevolved with ciliate” and “Evolved alone” panels). Most prey species exhibit similar variant frequency dynamics, such as selective sweeps that occur throughout the entire duration of the experiment. We have also observed non-trivial dynamics of variant clusters being maintained at a high frequency over long periods of time without these mutations getting lost or fixed (Fig. 4, “Coevolved with ciliate” and “Evolved alone” panels), suggesting previously described state of quasi-stable coexistence in bacterial populations (Good et al. 2017). The latter is clearly visible in the population 1 of *B. diminuta* in clades AC and AD, both undergoing their own sub-clonal expansions at different points in time. Other species exhibit similar clade coexistence, albeit with more stark difference in the clade frequencies. Similarly, selective sweeps occur in several study replicates, with coevolved *S. capsulata* population 3 displaying textbook successive sweeps. Successive hard selective sweeps are only observed in four cases; interestingly, all three coevolved *S. capsulata* populations, grown with ciliate, are among them, and the remaining one is a replicate of *C. testosteroni* grown without ciliate (Fig. 4, “Coevolved with ciliate” and “Evolved alone” panels). Furthermore, many populations show a pattern of late large scale diversification events, where many different clades start to appear in a quick succession halfway into the experiment (more than 400 days).

The number of clades consistently in both treatments increases over the duration of the experiment to the maximum of nine clades (linear regression coefficient of determination *R*^2^ = 0.75 and slope 0.0085 in coevolved populations, *R*^2^ = 0.69 and slope 0.0081 in evolved-alone populations). However, interestingly, the number of clades present at any given time in the experiment, after approximately 250 days, consistently stays at 3-5 clades (Supplementary Figs. 8 and 9) – suggesting active turnover of clades within a population as time goes on. This is likely explained by clonal interference between clades – however, since we do not observe consistent and repeated selective sweeps across the whole population, clonal interference dynamics are likely moderated by frequency dependent selection dynamics that sustain clade coexistence for longer periods of time (Maddamsetti, Lenski, and Barrick 2015; Good et al. 2017).

To quantitatively compare the inferred clonal structures between the experimental groups, in addition to Muller plots, we computed hierarchical representations in the form of strictly bifurcating phylogenetic trees (Fig. 4B, Supplementary Fig. 10, Supplementary Fig. 11), where the total branch length represents the number of days a clone existed since its emergence until either the end of the studied experiment or the final point in time the clone retains a frequency above 0.1 (provided no further sub-clones appear) – otherwise, the branch length is truncated. We computed commonly used tree metrics – sum, mean and variance of branch lengths, the number of leaves (corresponds to number of clones plus the background clone), maximum and average depths of the leaves, as well as Sackin, Colless and B1 indices of tree balance (Shao and Sokal 1990). Additionally, we computed RUI (robust-universal-interpretable) indices, as described in Noble and Verity 2023 – average effective outdegree, with and without accounting for branch lengths (^0^*D_N_* and ^1^*D_N_*, respectively), effective number of non-root nodes (^0^*D_S_*), effective number of branches, accounting for branch lengths (^1^*D_S_*), average branch count across the tree (^0^*D_L_*), effective number of maximally distant leaves (^1^*D_L_*), tree balance (^1^*J_N_* ), evenness of all branch lengths (^1^*J_S_*) and evenness of branch lengths across the tree (^1^*J_L_*). Using regression analysis, we first tested these metrics for statistically significant differences between the experimental groups (with predator and without predator) across all study replicates, using the species indicator as a covariate (ANCOVA; Rutherford 2001) – none of these measures differ significantly between the treatments (Supplementary Table 2).

We then compared the values of each tree index type between evolutionary histories for each species separately using ANOVA. Only one species – *B. diminuta* – exhibits a statistically significant difference between the evolutionary histories in only two of the metrics – the B1 and ^1^*D_N_* metrics (ANOVA, 95% CL; treatment group *n* = 3; B1 index *p* = 0.004, ^1^*D_N_ p* = 0.008). B1 metric measures tree balance – trees that have their leaves closer to the root on average are considered balanced, therefore this difference indicates that there is a greater imbalance observed in the structure of populations that evolved without ciliate. ^1^*D_N_* corresponds to average effective outdegree, accounting for branch lengths (Noble and Verity 2023). However, it must be noted that the tests on these two metrics would not be statistically significant either if multiple testing correction was applied (e.g., Bonferroni). For complete tables of within-species ANOVA results, see Supplementary Tables 3 and 4.

These results suggest that the presence of ciliate does not predictably affect the clonal structure of the studied prey populations, or, at the very least, the changes are subtle and remain largely hidden under shared evolutionary processes, such as adaptation to the growth medium. Instead, we observed the usual clonal interference and likely frequency dependent selection dynamics that universally characterize even simple microbial systems without predation (Maddamsetti, Lenski, and Barrick 2015).

### Mutation targets are recurrent both within and across species, and differ depending on the presence or absence of the predator

Lastly, we investigated the recurrence of non-synonymous variants identified by our genomic analyses to determine if there are patterns in recurrence and evidence of selection for mutations in specific genes. We defined the recurrence of a mutated gene as it being affected by at least one variant in at least two out of three replicates of a species, with and without the presence of the predator separately (Fig. 5, top panel). We also investigated the appearance times of the recurrent gene mutations, i.e., the appearance of the clade they belong in (Fig. 5, bottom panel) and found that the overall distribution of the timing of appearance, across all species together, is significantly differing between the evolutionary histories (two-sided Kolmogorov-Smirnov test p-value = 0.00003; Fig. 5, bottom left histogram). It appears that in the populations coevolving with the predator, more recurrently targeted genes are affected late in the experiment (*>* 600 days) than in populations evolving alone, moreover, only the coevolving populations show recurrently affected genes appearing within the first 100 days of the experiment.

**Figure 4:**
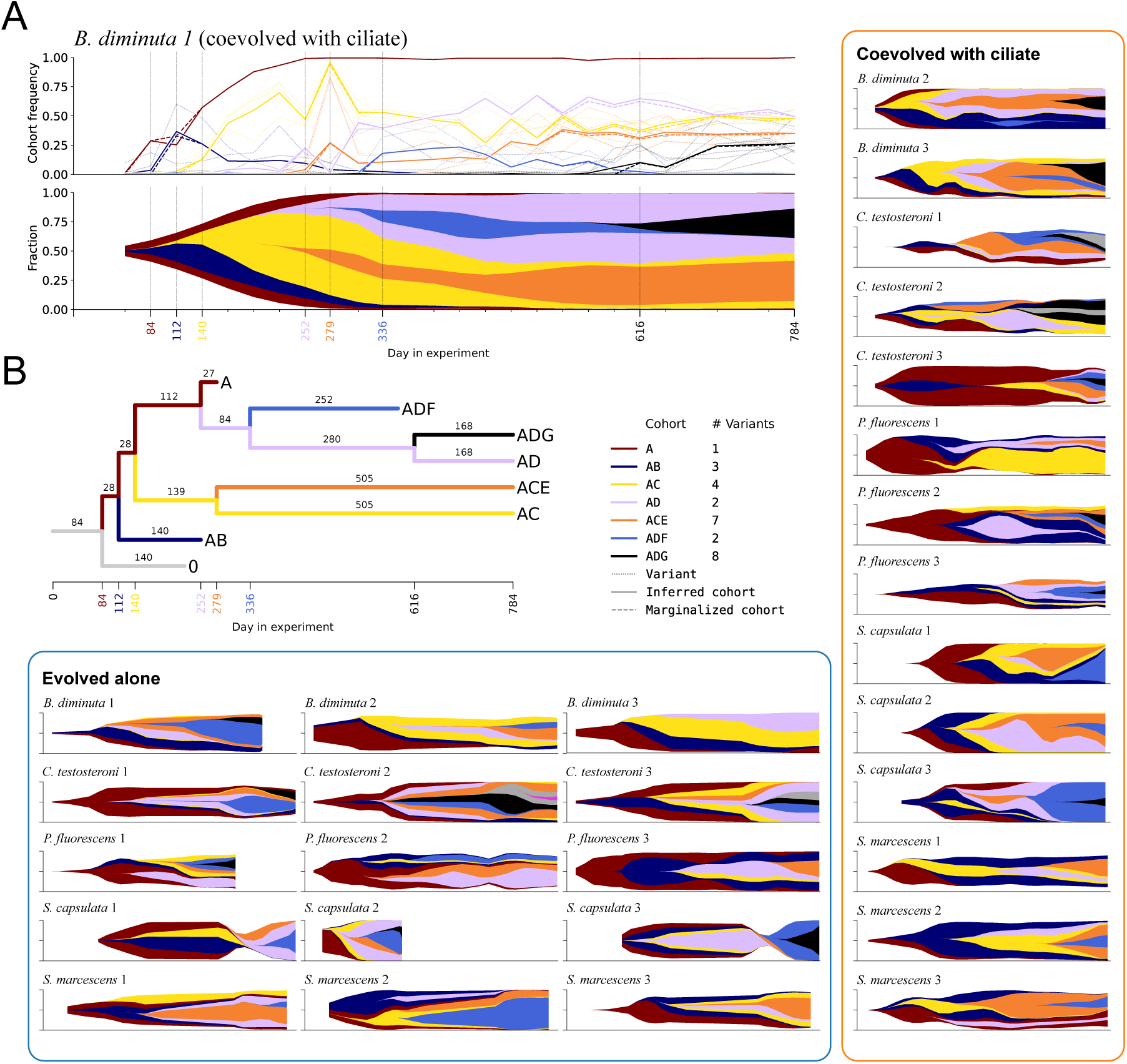
A. Clonal structure evolution in *B. diminuta* under predation (replicate population 1) in terms of variant frequency trajectories over the course of the long-term experiment. Trajectories are first clustered into cohorts and then assigned to a likely position in the clonal structure (see Methods) and represented as a Muller plot. Vertical dashed bars in the trajectory and the Muller plot mark the appearance of the clone (cohort trajectory exceeding the emergence threshold, see Methods). **B** A corresponding phylogenetic tree representation of the population structure of *B. diminuta* under predation (replicate population 1). The time of the appearance of each clone is color-coded to correspond to the clone information in the legend and the variant trajectories with shaded areas in the trajectory and Muller plots, respectively. The branch labels in the tree denote the time the clone remains above a survival threshold (provided no further sub-clones emerge; see Methods). A non-trivial long-term coexistence of the inferred clades is clearly seen in the Muller plot and the phylogenetic tree. **Evolved alone** and **Coevolved with ciliate** boxes: Muller plot showcase for the remaining 29 populations evolving under and without predation, respectively. There is significant variability both between species and between replicate populations of the same species. Long-term coexistence of clades is a recurring theme, along with examples of selective sweeps. Populations with truncated Muller plots represent populations where a portion of time series data was lost to contamination during laboratory procedures.

**Figure 5:**
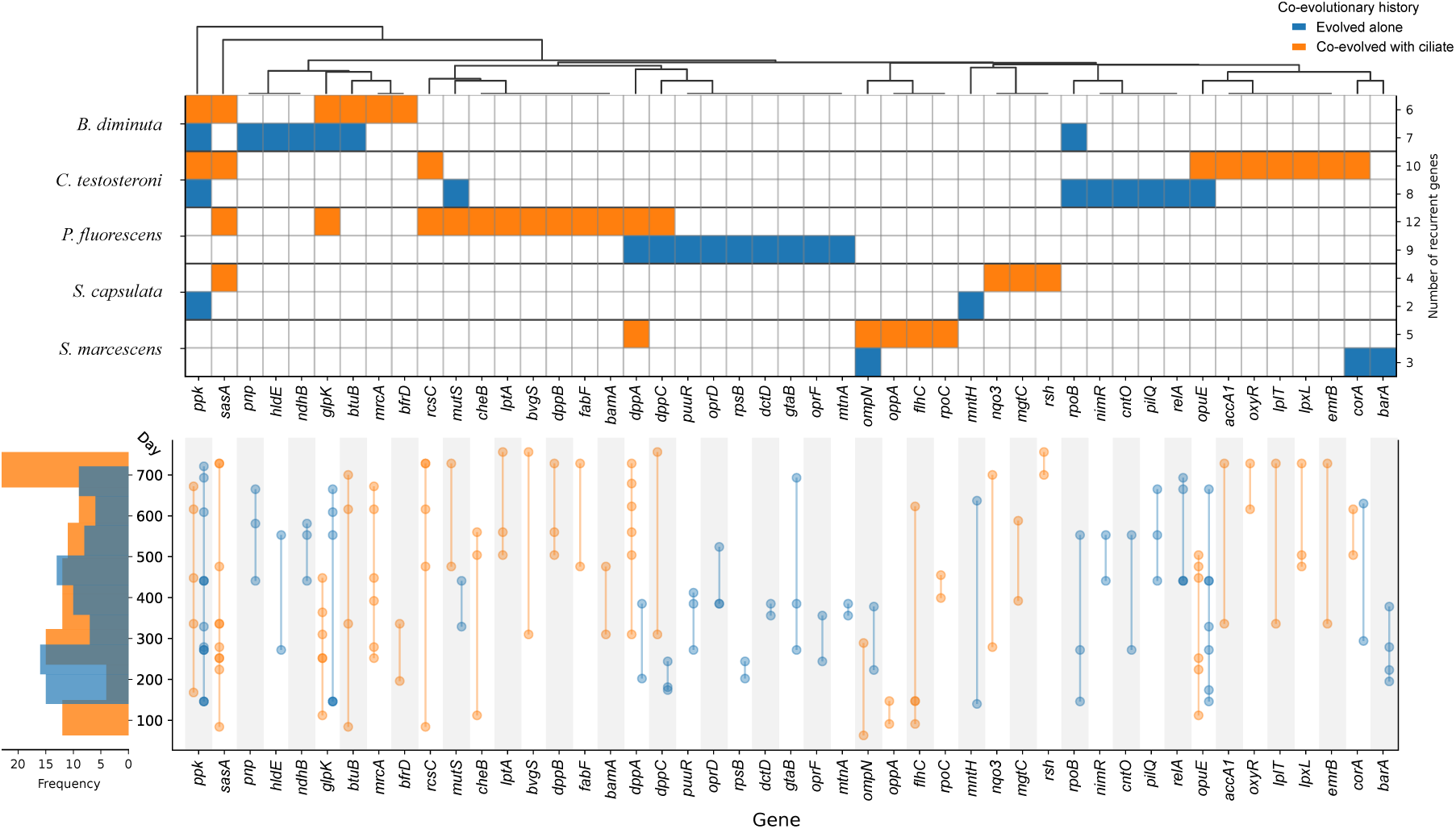
Top panel: non-synonymous variant hit recurrency across the experiments in known genes. Columns (i.e., genes) are hierarchically clustered according to the pairwise Hamming distance. Populations without ciliate (blue rows) have 17 recurrent genes containing variants that do not occur in populations grown with the ciliate (orange rows). Conversely, the set of genes that are hit only in populations with ciliate is larger (21 genes). Bottom panel: variant timing of appearance (based on cohort appearance to which a variant belongs to). Darker circles indicate that there are multiple unique study replicates where a variant in a given gene appeared at the same time. Marginal counts of variant appearance timing are shown as a histogram.

We computed gene hit indicator data (1 if a non-synonymous variant has been detected in a gene at least once, 0 otherwise) for all genes. Then, using ANCOVA with species indicator as a covariate, we tested whether the means of hit indicators differ between evolutionary histories (95% CL, treatment group *n* = 15 per gene). Eleven genes – *barA*, *cheB*, *dppB*, *flhC*, *lptA*, *puuR*, *rcsC*, *relA*, *rpoB*, *rpoC* and *sasA* – have been identified as having a significantly differing recurrence between the experimental groups, i.e., growth with ciliate versus without ciliate (Fig. 5, Supplementary Table 5).

Out of these, seven – *cheB*, *dppB*, *flhC*, *rpoC*, *lptA*, *rcsC* and *sasA* – are recurrent only in the coevolved populations. Gene *cheB* (also known as *wspF* ) is recurrent in all three coevolved *P. fluorescens* populations and has been previously reported to be also recurrently targeted in a dual-stressor (antibiotic and ciliate) system with *P. fluorescens* as prey (Hiltunen et al. 2018). Periplasmic dipeptide transport gene *dppB* is part of the dipeptide permease complex and has been associated with bacterial growth (Abouhamad and Manson 1994), and is only affected in all three coevolved *P. fluorescens* populations. Genes *flhC* and *rpoC*, recurring only in coevolved *S. marcescens* populations (all three and two out of three, respectively), are associated with biofilm formation (Sule et al. 2011) and RNA polymerase subunit *β*’ (Finn et al. 2000), respectively. Lipopolysaccharide transport protein A gene *lptA* is responsible for cell structure integrity (Suits et al. 2008), and is recurrently affected only in coevolved populations of *P. fluorescens* populations (two out of three). Hybrid sensor kinase gene *rcsC* has been identified as an important part of *E. coli* biofilm formation (Ferrìeres and Clarke 2003) and is recurrently affected in all coevolved replicates of both *C. testosteroni* and *P. fluorescens*. The gene encoding the adaptive sensory kinase *sasA*, recurrently affected in coevolved-only populations across all studied species except *S. marcescens* (two out of three coevolved populations of *B. diminuta*, *C. testosteroni*, *P. fluorescens* and all three coevolved populations of *S. capsulata*), is purported to be a core part of the circadian rhythm control mechanism in cyanobacteria (Iwasaki et al. 2000), though to our knowledge, its exact physiological role in other bacteria is not known.

In contrast, four genes – *barA*, *puuR*, *relA* and *rpoB* – are recurrent only in evolved-alone populations. Variants in the bacterial adaptive response gene *barA*, coding for signal transduction histidine-protein kinase (Sahu et al. 2003), are detected in all three no-predator populations of *S. marcescens*, but not in any other species’ experiments. Gene *puuR* is affected in all three evolving-alone populations of *P. fluorescens* and is associated with putrescine utilization (Kurihara et al. 2005), which promotes biofilm formation in some species (Liu et al. 2022). Genes *relA* and *rpoB* are associated with biofilms (Boehm et al. 2009) and various RNA polymerase *β* functions (Mollet, Drancourt, and Raoult 1997), respectively; both are recurrent in *C. testosteroni* evolved-alone populations (all three and two out of three, respectively) and *rpoB* is also recurrent in two out of three *B. diminuta* evolved-alone populations, thus likely not related to defense against the predator.

The majority of remaining recurrent genes, albeit with no significance in recurrence between the evolutionary histories, are limited to recurrence in singular species (Fig. 5, top panel). Out of these genes, the ones that are recurrent in coevolving populations only are mostly responsible for various metabolic functions, cell wall formation and maintenance, though some are related to biofilm formation, which is expected given bacteria propensity to form biofilm in response to external stress, such as predation, in addition to simply being a viable strategy to adapt to the environment (Kimberly K Jefferson 2004). Many genes that recur in both experimental treatments are related to various metabolic functions and are likely responsible for adaptation to the growth medium rather than defense. In total, only eight genes – *ppk*, *sasA*, *glpK*, *rcsC*, *mutS*, *dppA*, *rpoB* and *corA* – are recurrently affected in more than one of the species, regardless of the evolutionary history.

## Discussion

With our long-term experimental evolution experiment, we aimed to study whether a strong selective pressure, i.e., predation, would significantly change the clonal dynamics and the outcome of molecular adaptation in the prey bacterium. In most of our study species, we observed clear signs of phenotypic adaptation, where coevolved bacterial prey populations grow substantially better in the presence of ciliate compared to control populations. Moreover, putative mutation counts are higher in coevolved populations and recurrently mutated gene sets are different between the experimental groups – thus, both phenotypic and genomic data support treatment-specific evolution. Given the fact that we found more recurrent mutation targets to appear both earlier and later in coevolved populations than in the evolved-alone populations (Fig. 5, bottom panel), we speculate that the ciliate forces the prey to evolve defenses quickly, and the pressure by the ciliate is kept up throughout the entire experiment. The latter point is supported by our expectation that the ciliate coevolves alongside the prey – newly evolved prey defenses cause ciliate to evolve counter-defenses, to which the prey has to again adapt, and so on. Population sizes of predator and prey in coevolved populations, as expected, fluctuate in opposition to one another and are negatively correlated (linear regression *R*^2^ = −0.42, slope −1.78, Supplementary Fig. 12). However, surprisingly, despite our observance of rich and non-trivial clonal structure dynamics, we cannot detect a strong signal of predation within the clonal structures of our study populations. We did not observe systematic changes in clonal structure evolution pattern, such as an increased number of selective sweeps, like we hypothesized that it could be one reflection of ciliate’s selective pressure on the prey. Instead, we find that the observed molecular evolutionary dynamics of bacteria subject to predation are indistinguishable from those of bacteria evolving alone – characterized by known processes, such constant dynamic emergence of adaptive clades, and coexistence and competitive dynamics between them (Good et al. 2017).

What we observed in this study may be a case of “emergent simplicity” – complex and unpredictable genomic evolution is ultimately resulting in predictable phenotypic changes (Lässig, Mustonen, and Walczak 2017). There may be many metabolic pathways through which prey bacterium adapt to their immediate environment (both the growth medium and the ciliate), changing their mode of growth in a predictable manner, but on a genomic level, prey population clonal dynamics may be largely driven by the aforementioned universal evolutionary processes, much like in previous studies on pairwise bacterial community evolution (Meroz et al. 2024). Moreover, it has been suggested that the repeatability of evolution may not necessarily be mainly driven by strong selective pressure (Bailey et al. 2017). Partly repeatable evolution in terms of clade dynamics has been described in influenza (Strelkowa and Lässig 2012; Łuksza and Lässig 2014), where strong clonal interference drives successful clades to constantly emerge victorious over the other clades in the population – that is predominantly not the case in our clade trees, and instead we observed constant competition and coexistence of clades. An interesting thing to note is that we observe rapid changes in clonal diversity that occur rather late – in some cases, even in the final 100 days in our 800 day-long experiment (Fig. 5) – so it would be interesting to see whether there are any significant changes in clonal structure over much longer timescales, though that can be, unfortunately, time and cost prohibitive.

There might be several explanations for the lack of clearly observable predator signal in clade dynamics in our study: 1) our data resolution might be insufficient to observe subtle changes in between genomic sampling (every four weeks in our study); 2) the predator in our study, *Tetrahymena thermophila*, is a generalist predator, and therefore may not exert a strong enough selective pressure to influence clade dynamics; and 3) frequency dependent selection, which prevents a quick competitive exclusion of the prey clades (Maddamsetti, Lenski, and Barrick 2015), unlike in influenza evolution. Moreover, within each population, there is a constant maintenance of several clades coexisting at the same time above threshold frequencies (Supplementary Fig. 8).

Also, we recognize that our statistical power (three replicates per species, per treatment) might be a limiting factor in detecting subtle clonal dynamics. To fully disentangle the clonal dynamics of coevolving prey, we believe that a larger number of phenotypic measurements (than 10 clones per sampling time point in our study) would be needed from every sampling time point to account for the ever-increasing clonal diversity of the populations – growth profiles of the limited number of isolates taken from a time point where one clone dominates might be hiding subtle effects of clones that are present at lower frequencies. Additionally, since the first time point of isolate sampling represents populations that have already (co-)evolved for at least a month, we cannot compare early prey growth dynamics between the treatments. Furthermore, sampling the populations for genomic data every four weeks might be too coarse to account for rapid eco-evolutionary dynamics, such as overall prey and predator population size fluctuations during one-week transfer cycles (see Methods), though it is not clear whether increasing the overall time resolution with respect to the genomic data sampling alone would result in different conclusions. Similarly, due to the limited number of biological replicates per species in our experiment, it is difficult to make definitive conclusions regarding the significance of mutation gene target recurrence. While the patterns of recurrence do not seem to arise by chance, it is unclear to what extent each putative recurrent mutation drives the adaptation, and if they represent adaptation to the growth medium, the ciliate, or both.

For future studies, we believe that long-term coevolution experiments need carefully designed linking between genomic and phenotypic data tracks in order to disentangle the impact of strong environmental pressure (i.e., predation) from other evolutionary processes. Given that our fitness measure for the studied prey is only a proxy derived directly from growth curves (i.e., growth relative to the ancestor in terms of AUC; Fig. 2), phenotypic analyses could be expanded to include competition assays in order to make stronger conclusions regarding phenotypic variation and treatment differences. The dynamics of clones, represented by variant cohorts of SNPs and indels, seem to quickly become so complex that differentiating predation pressure from other evolutionary processes remains an outstanding challenge. Furthermore, it would be interesting to study the coevolution of the predator as well and contrast the evolution of prey defense to evolution of predator offense – for that, one would need to produce equally rich genomic data from the predator and investigate clonal dynamics and gene recurrence in a similar fashion, though it is expected that the prey would experience greater evolutionary changes than the predator. Nevertheless, studying predator coevolution in terms of predator population clade dynamics might yield interesting novel insights.

While it is still surprising that in this study we do not observe differences in clade dynamics between our evolutionary treatments, given the strong phenotypic and genomic effects from predation, we believe that the predator signals, while being detectable, remain masked within the clade dynamics by pervasive prey clade interactions, stable coexistence and frequency dependent selection that maintains both undefended and defended types within the prey populations. We believe that our study further reinforces the non-triviality of clonal dynamics even in the presence of a strong selective pressure despite easily detectable phenotypic differences and also provides new methodology to represent and investigate the clonal structure of a population solely through genomic time series data.

## Materials and Methods

### Experimental setup

The experiments described in this article are a five-species subset of long-term continuation of seven species (co-)evolution experiments described in Cairns et al. 2020a, where bacterial prey species *Brevundimonas diminuta* HAMBI 18, *Comamonas testosteroni* HAMBI 403, *Pseudomonas fluorescens* SBW25, *Sphingomonas capsulata* HAMBI 103, *Serratia marcescens* ATCC 13880, *Escherichia coli* ATCC 11303 and *Janthinobacterium lividum* HAMBI 1919 were preyed upon by a ciliate *Tetrahymena thermophila* CCAP 1630/1U. While we continued the evolutionary experiments and collected genomic and phenotypic data for all seven species, *E. coli* and *J. lividum* were omitted from this study for data completeness considerations: the majority of *E. coli* genomic data was of poor alignment quality (indicating sample contamination), and we could not revive *J. lividum* stocks for phenotypic sampling. The laboratory procedures of the continued experiments were identical to the original experiment, but here we recapitulate the key details.

The evolutionary experiment was started using an aliquot (20 µl) of a 48 h bacterial culture, started from a single colony and ∼10 000 ciliate cells (∼1700 cells ml^−1^) from an axenic culture. Each bacterial strain was cultured both alone and together with the ciliate predator (three replicates each) in batch cultures of 20 ml glass vials containing 6 ml of 5% King’s B (KB) medium, with 1% weekly transfer to fresh medium. The KB medium recipe (concentrations in 100% medium) is as follows: 20 g Peptone number 3 and 10 ml of 85% glycerol in 1 litre of dH_2_O. During the experiment, cultures were kept at 28°C (±0.1°C) with shaking at 50 r.p.m. Every four transfers (i.e., 28 days), population samples were freeze-stored (frozen in 28% glycerol and kept at -80°C) and bacterial density was estimated based on optical density at 600 nm wavelength, and ciliate density was estimated based on light microscopy counts (5 × 0.5 µL droplets) according to previously established protocols (Hiltunen et al. 2018; Cairns et al. 2016). The ciliate was cultured axenically, i.e., without other bacteria prior to co-culture to prevent bacterial contaminants. Predator cells were made axenic 1-2 times per year by transferring coevolving cultures to 50 ml of 20% R2A (Reasoner’s 2A agar; commercial product by Labema, Helsinki, Finland) medium, then to antibiotic cocktail in 6 ml PPY (Proteose Peptone Yeast extract medium), and finally to 50 ml PPY for growth. PPY recipe: 20 g of proteose peptone and 2.5 g of yeast extract in 1 l of deionized water. Axenic predator cells were stored in liquid nitrogen.

We have collected data for populations of the seven species coevolving with *T. thermophila* for a period of ∼1400 days. Originally, the experiment was designed to follow co-culture coevolution in a similar fashion to long-term E. coli evolution in Good et al. 2017, and to make comparisons of coevolutionary outcomes between different prey species. In that setup it is possible to study prey defense and predator counter-attack evolution (Hiltunen and Becks 2014), which does not necessitate prey control lines (bacteria evolving alone). However, ∼600 days into the coevolutionary experiment, we also decided to start control lines in order to be able to investigate the effect of predation on prey molecular dynamics. The experiments in both lines finished at the same time, but because of the late start of the control line, it corresponds to a shorter period of evolution. Therefore, because this present study largely focuses on molecular evolution in prey, we only investigated the data corresponding to the duration of the experiment common to both evolutionary lines (∼800 days). In total, 42 bacterial populations (7 species, 2 evolutionary lines, 3 replicates each) have undergone at least 800 days of evolution (∼1400 days for coevolving prey and ∼800 days for prey evolving alone). Considering the prey genomic sampling (from freeze-stored whole-population samples) resolution of ∼1400/28 for coevolving populations and ∼800/28 for populations evolving alone, we sampled and sequenced 1694 genomic samples corresponding to the seven study species, three replicates each for both evolutionary lines. Out of these, for this study, we analysed 30 bacterial populations (5 species, 2 evolutionary lines, 3 replicates each) that have undergone at least 800 days of evolution and are represented by 1221 time-resolved genomic samples.

### Genomic data and variant calling

Bacterial DNA was extracted from freeze-stored whole-population samples (corresponding to intervals of 28 days in our evolutionary experiments) using the DNeasy Blood & Tissue 96 Kit (Qiagen) according to the manufacturer’s protocol. DNA concentration was measured with the HS assay kit using the Qubit 556 3.0 fluorometer (Thermo Fisher Scientific, Waltham, MA, United States). Libraries for each single-species bacterial population sample DNA were prepared with Nextera DNA Flex and bulk sequencing performed using short-read next–generation sequencing (NGS) equipment Illumina NovaSeq 6000 (read length 150). To obtain the reference genome sequences of our study species, we sent our ancestral (i.e., unevolved) samples to the PacBio HiFi long-read sequencing laboratory (URL: https://www.pacb.com) for long-read sequencing and assembly of the genome.

After assessing the read quality metrics of each prey population sample, genetic variants were obtained by following this read processing, variant calling and annotation workflow: Illumina DNA adapters trimmed with *cutadapt* (version v3.5; NBIS) (Martin 2011), reads aligned to reference sequences with *BWA* (version 0.7.17; Li and Durbin) (Li and Durbin 2009), read duplicates marked with *Picard* (version 2.27.5; Broad Institute, URL: https://broadinstitute.github.io/picard/), variants called in a multi-sample mode and subsequently filtered with *GATK Mutect2* (version 4.3.0.0; Broad Institute) (Benjamin et al. 2019), and finally annotated with *snpEff* (version 5.1d; Pablo Cingolani) (Cingolani et al. 2012).

Since we have also sequenced samples for populations that coevolved with the predator beyond 800 days (up to ∼1400 days; see Experimental setup section in Methods), all samples have been used to call the variants. However, because populations evolving alone (controls) were limited to a 800 day period, the variant data corresponding to samples taken later than 800 days into the experiment were not included in any downstream analyses of this study (see Supplementary Information for sample curation and tool usage details).

### Phenotypic data

From each bacterial population in our study, in addition to four ancestral (day 0) isolates, ten isolates were taken at two time points in the experiment. The first sampling, early in the experiment, is done at days 135 (*S. marcescens*) and 64 (rest of the study species) from the evolved-alone populations; and at day 43 (all study species) from the coevolved populations. The second sampling, late in the experiment, is done at days 723 (*S. marcescens*) and 652 (rest of the study species) from the evolved-alone populations; and at day 631 (all study species) from the coevolved populations.

The difference in sampling timing between coevolved and evolved-alone evolutionary lines was because evolved-alone lines were started later than coevolved ones (see Experimental setup section in Methods), resulting in slightly different freeze-storing schedules. Additionally, the evolved-alone *S. marcescens* lines were started before the lines of all other study species, but all experiments were sampled at the same date; therefore, evolved-alone *S. marcescens* correspond to a slightly longer period of evolution when both (early and late) samplings were done for either evolutionary line.

The isolates were then grown in three kinds of media: 5% King’s Broth (KB) medium (without the presence of ciliate), predator medium (with ciliate) and salt stress medium (1% w/v potassium chloride KCl; without ciliate). Growth curves were extracted by measuring the optical density of the grown bacterial colony every 20 minutes using the Agilent LogPhase 600 nm device. For each isolate, optical density measurements were replicated three times in total. In total, there were 5 × 2 × 3 × 2 × 10 × 3 = 1800 growth curves (5 species, 2 evolutionary treatments, 3 replicate populations, 2 time points, 10 isolates, 3 technical replicates). Due to some inherent variability in these technical replicate measurements, we computed their mean at each time point and used the resulting curve as our representative growth curve of that isolate, totaling in 600 growth curves that were analysed further. Additionally, there were four ancestral strain isolates taken for each of the five species and for each of the three growth media, and measurement was replicated four times, totaling 5 × 3 × 4 × 4 = 240 ancestral growth curves.

For each growth curve *N* (*t*)*, t* = 1*, . . ., t_max_* we computed its per-capita growth rate curve *ρ*(*t*) as:

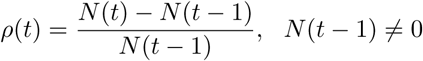

If *N* (*t* − 1) = 0, we set *ρ*(*t*) = 0. Then *ρ_max_* is the maximal value of *ρ*(*t*) and *t_ρ__max_* is the time when *ρ* attains its maximum, i.e., *ρ*(*t_ρ__max_* ) = *ρ_max_*. We compute yield *Y* as the final value of the growth curve *N* (*t*), i.e., *Y* = *N* (*t_max_*), and the area under the curve (AUC) as 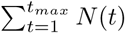.

To compute ratios of AUC between isolates of either evolutionary treatment versus their ancestral counterparts, we computed the mean at each time point of the four technical replicates of ancestral growth per species, per growth medium (control, salt stress and predator media), resulting in 60 ancestral mean growth curves. Then, for each species *s*, the technical measurement mean AUC of each isolate *i* from populations of either evolutionary treatment (denoted *E*) were divided by the corresponding species ancestral mean AUC (denoted *A*) in the relevant growth medium *m*:

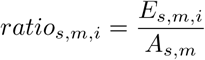

### Linear mixed models

To model the relative fitness values using evolutionary history, species identity, sampling timing (fixed effects) and replicate population identifier (random effect), we used *lme4* (Bates et al. 2015; version 1.1-38) package for *R* programming language (R Core Team 2025; version 4.5.2), as implemented in *RStudio* (Posit team 2025; version 2025.09.2, build 418). Our base model was specified as “Ratio ∼ History + Species + Timing + (1|Replicate)”, where “Ratio” is the relative fitness, “History” is the evolutionary history, “Species” is the species identity, “Timing” is the isolate sampling timing, and “Replicate” is the replicate population identifier, specific to each species in either evolutionary history. The interaction model was specified as “Ratio ∼ History + Species + Timing + History:Species + History:Timing + Species:Timing + (1|Replicate)”. The *R*^2^ values, reported throughout the main text, were computed from model-fitted values and observed fitness values. Proportion of variance explained by each of the main terms (Fig. 2) was computed using *partR2* package (Stoffel, Nakagawa, and Schielzeth 2021; version 0.9.2).

To determine the significance of each of the three main fixed linear mixed model terms, we used a log-likelihood ratio test as implemented in *R*’s *anova* function, where the two arguments are: 1) the base linear mixed model, and 2) base model without the tested term. To determine the significance of each interaction term, the arguments are: 1) the base linear mixed model, and 2) base model plus the tested interaction.

### Random forest classifier

To assess the classification accuracy in classifying phenotypic growth profiles (growth curves from all three growth media combined) according to species, evolutionary history, replicate and sampling time, we used scikit-learn (Pedregosa et al. 2011) (version 1.4.2) Python (version 3.10.14) implementation of the random forest classifier to train 100 random forest models and averaged the resulting confusion matrices representing the test dataset over all model instances. The final averaged confusion matrix (Fig. 3) is an element-wise sum of all produced confusion matrices, divided by the number of model instances, i.e., 100. Before learning each model, phenotypic growth profile data was randomly split in half for training and testing datasets in a stratified fashion, i.e., keeping the proportion of each unique label equal across both datasets. For result repeatability, each train-test split and random forest model instance were generated with a Numpy (Harris et al. 2020) (version 1.26.4) random seed *s* = 0*, . . .,* 99 for each of the 100 runs.

### Clonal structure inference from mutation cohorts

Variant trajectories have been assigned to variant cohorts (clusters) using read counts of reference and alternate alleles with PyClone-VI (Gillis and Roth 2020). Though any other similar clustering software would likely be sufficient for this task, we chose PyClone-VI for its acknowledged status in bioinformatics research and ease of use. Since this software is originally intended for human cancer genomics and the genomes under our study are haploid, we set the required parameters for major and minor copy numbers to one and zero respectively for every variant within the genomic time series data of any given population from our experiments. For each replicate separately, using 15 for an initial guess for the number of clusters with 100 restarts of the inference algorithm, we repeated the inference ten times, each with a different random seed (seed values 1, 2, …, 10), and selected the model with the highest final evidence lower bound (ELBO) value, as described in the PyClone-VI manuscript (Gillis and Roth 2020). The final number of inferred clusters varied from four to nine – however, we note that repeated inferences with different seed values may yield slightly different variant trajectory clusters and their numbers.

Given the computed variant cohorts of a prey population, we infer its clonal structure by following two clonality rules: 1) for two competing clones to co-exist, the sum of their frequencies cannot exceed 1 at any time point, and 2) the sum of the frequency of every sub-clone cannot exceed the frequency of the clone they belong to. The hierarchy of the clonal structure is represented by a tree, where each node represents a mutation cohort. As the cohorts “emerge” over time, i.e., their frequency exceeds a chosen threshold at a given time point (0.1 by default; some replicates needed higher thresholds, e.g., 0.15, to resolve obviously wrong hierarchical assignments due to multiple cohorts emerging at the same time with a similar frequency, close to the threshold), they are added as new nodes to the growing trees. After the first cohort has emerged and has been placed under the root node of the tree, every subsequent emerging cohort creates new trees, each representing one of the possible placements of the new node within each tree constructed before. Therefore, the total number of such possible trees that include every mutational cohort is the factorial of the number of inferred cohorts. The order of the cohort emergence is known, except in cases where two or more cohorts exceed the emergence threshold at the same time, doubling the number of possible trees with each additional emerging cohort at such a time point – because of that, we specify the order manually to prevent possibly degenerate cases due to incorrect clonality role assignment and to limit computational cost.

Next, having a set of complete clonal hierarchy trees for a given population, we use the mutational cohort frequencies *Q* to calculate the allele frequencies *F* that correspond to the inferred clonal structure at a given time *t*. Given a clone *c* and using *x* ⊂ *c* to denote that clone *x* is a sub-clone of *c*, we define the allele fraction as:

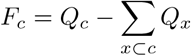

Since the originally computed mutation cohort frequencies are inherently noisy because of the variant trajectory variation within trajectory clusters, we use the newly calculated allele frequencies to marginalize the cohort frequencies, i.e., adjust their values per time point so that they fully agree with the two clonality rules. That is done by iteratively subtracting a small value (0.1% of the absolute *F_c_* value) from the frequencies of alleles under consideration until their respective sum is either less than 1 or less than the frequency of the clone they are sub-clones of, respective to the placement of the considered allele in the previously inferred tree structure. This way, we evaluate the consistency of the inferred clonal hierarchy both quantitatively and visually.

Lastly, we select a tree with the lowest total marginalization cost to be the representative clonal structure of a given replicate. Since the marginalization of every possible tree is computationally time-prohibitive, we first select a subset of most likely tree configurations by implementing an arbitrary scoring function which corresponds to the two clonality rules; when considering a node that represents a cohort that emerged at time *t*, the score is the number of future time points *t_f_ > t* where neither of the rules are violated, allowing up to 0.01 of absolute error to account for the potential noise in the frequency data. While the minimum score usually represents low-cost solutions, they may not be the lowest possible ones, and an objective solution other than the marginalization is not known. To help mitigate this, we marginalize 50 best scoring clonal structures and choose the one with the minimal cost as our representative structure. Inferred clonal structures presented in this article are intended as examples of automated, yet realistic, solutions and not objectively best ones – it would likely be possible to find marginally lower-cost solutions.

We note that multiple rule-consistent and high scoring solutions are sometimes possible, especially when low-frequency cohorts emerge late in a population with multiple competing clones present at sufficiently high frequencies. Additionally, some cohorts never reach our emergence threshold, and we remove these cohorts altogether. Nevertheless, this method could be extended to potentially handle every possible configuration of the clonal structure tree in a more efficient way.

## Data availability

Raw sequence data (bam files) will be deposited in the EMBL ENA. Accession codes and links to be added upon acceptance of the manuscript. All code and pre-processed data needed to reproduce the downstream analyses and figures will be made available upon publication via GitHub: https://github.com/dovydask/clonal_structure.

## Author contributions

J.C. and T.H. designed the long-term evolution experiment. J.C. and V.P. performed the long-term evolution experiment. J.H. performed phenotypic measurements. D.K. performed sequence data bioinformatics, downstream analyses and algorithm design. D.K., J.C., T.H., L.B. and V.M. were responsible for interpreting results. D.K. and J.C. wrote the original paper draft, with contributions from all authors. J.C. T.H., L.B. and V.M. supervised the work. All authors approved the final version of the paper.

## Acknowledgments

We thank Liisa Ruusulehto, Paula Typpö, Roosa Jokela, Eeva Vakkari, Jutta Kasurinen, Iina Hepolehto, Tinja Hyvönen for their laboratory procedure work on the long-term experiment. We thank the WGGC Core Facility Cologne for the preparation and sequencing of our samples. We also thank Anthony Sun and Mikko Kivikoski for their helpful insights and comments on algorithmic and bioinformatic procedures.

