## Supplementary Information for "Universal bacterial clade dynamics dominate under predation despite altered phenotypes and mutation targets"

February 2025

#### Sequencing data bioinformatics

##### Raw data

Sample metadata is provided in *sequencing_sample_information.csv* file in our GitHub code repository. Samples with "PS" prefix denote prey populations evolved with the ciliate ("Predator Selection"), and samples with "NP" denote prey evolved without the predator ("No Predator"). Samples with a low ($< 50\%$) reference sequence alignment have been excluded, as well as ones with suspected lab errors in measurement or unexpected culture conditions ("unreliable population size" column). "Selected for clonal inference" column denotes the samples from which variant frequencies have been extracted after multi-sample variant calling and used to determine mutation cohorts (through PyClone-VI).

All bioinformatics pipeline tools were used using default tool-specific parameters unless otherwise specified.

##### Adapter trimming

Adapter sequences trimmed with *cutadapt* (version v3.5); bases of quality $< 28$ removed before adapter trimming. Minimum read length 30 and minimum overlap 10.

Read 1:
AGATCGGAAGAGCACACGTCTGAACTCCAGTCA
AGATCGGAAGAGCACACGTCTGAAC
TGGAATTCTCGGGTGCCAAGG
AGATCGGAAGAGCACACGTCT
CTGTCTCTTATACACATCT
AGATGTGTATAAGAGACAG

Read 2: AGATCGGAAGAGCGTCGTGTAGGGAAAGAGTGT
AGATCGGAAGAGCGTCGTGTAGGGA
TGGAATTCTCGGGTGCCAAGG
AGATCGGAAGAGCACACGTCT
CTGTCTCTTATACACATCT
AGATGTGTATAAGAGACAG

##### Alignment to the reference sequence

Paired read alignment was performed with *BWA* using *bwa mem* command. The resulting sam file was converted into a bam file using *samtools view*, sorted using *samtools sort* and indexed using *samtools index*. Duplicates marked using *picard MarkDuplicates*, read groups added using *picard AddOrReplaceReadGroups*, reindexed with *samtools index* and alignment statistics collected with *picard CollectWgsMetrics* and *picard CollectAlignmentSummaryMetrics*. Population-specific samples of the same date (from multiple sequencing lanes) were merged into one bam file using *samtools merge*.

#### Variant calling

Multi-sample variant calling per study population was done using GATK (version 4.3) using *gatk mutect2* command with the first sample (by date) used as a normal and the rest of samples (see sequencing data information file) were provided as input files. Variants filtered using *gatk FilterMutectCalls* in microbial mode. Only the SNPs and Indels were selected using *gatk SelectVariants* and all variants without a PASS flag were excluded.

#### Variant annotation and downstream analyses

Reference genomes were annotated using *Prokka* (version 1.14.6) and genomic variants were annotated using *snpEff* (version 5.1d). All subsequent allele frequency extractions and analyses were done with Python programming language.

#### PyClone-VI usage

Variant frequency data was formatted according to PyClone-VI requirements – in order to make the software applicable to haploid bacterial DNA data, major, minor and normal copy numbers set to 1, 0 and 1 respectively. The resulting data were used as an input PyClone-VI, using 15 fitting clusters (-c parameter), 100 random restarts for variational inference (-r parameter). All other parameters were left as default. We did ten runs with a seed (values 1, 2, ..., 10) for result replicability and selected the representative run with the highest evidence-lower bound (ELBO) value.

Samples that were excluded from clonal inference after mutation cohort inference step due to artefactual properties (such as their variant frequencies being all zero).

- PS_Ct_r02_2016_05_04_L001
- NP_Bd_r03_2018_03_06_L001
- NP_Ct_r01_2017_08_21_L001
- NP_Ct_r01_2018_03_06_L001
- NP_Ct_r02_2018_03_06_L001
- NP_Ct_r02_2018_05_01_L001
- NP_Ct_r03_2018_03_06_L001
- NP_Ct_r03_2018_04_03_L001
- NP_Ct_r03_2018_05_01_L001
- NP_Pf_r01_2018_07_24_L002
- NP_Sc_r01_2017_12_12_L001
- NP_Sc_r03_2018_03_06_L001

### Supplementary Figures

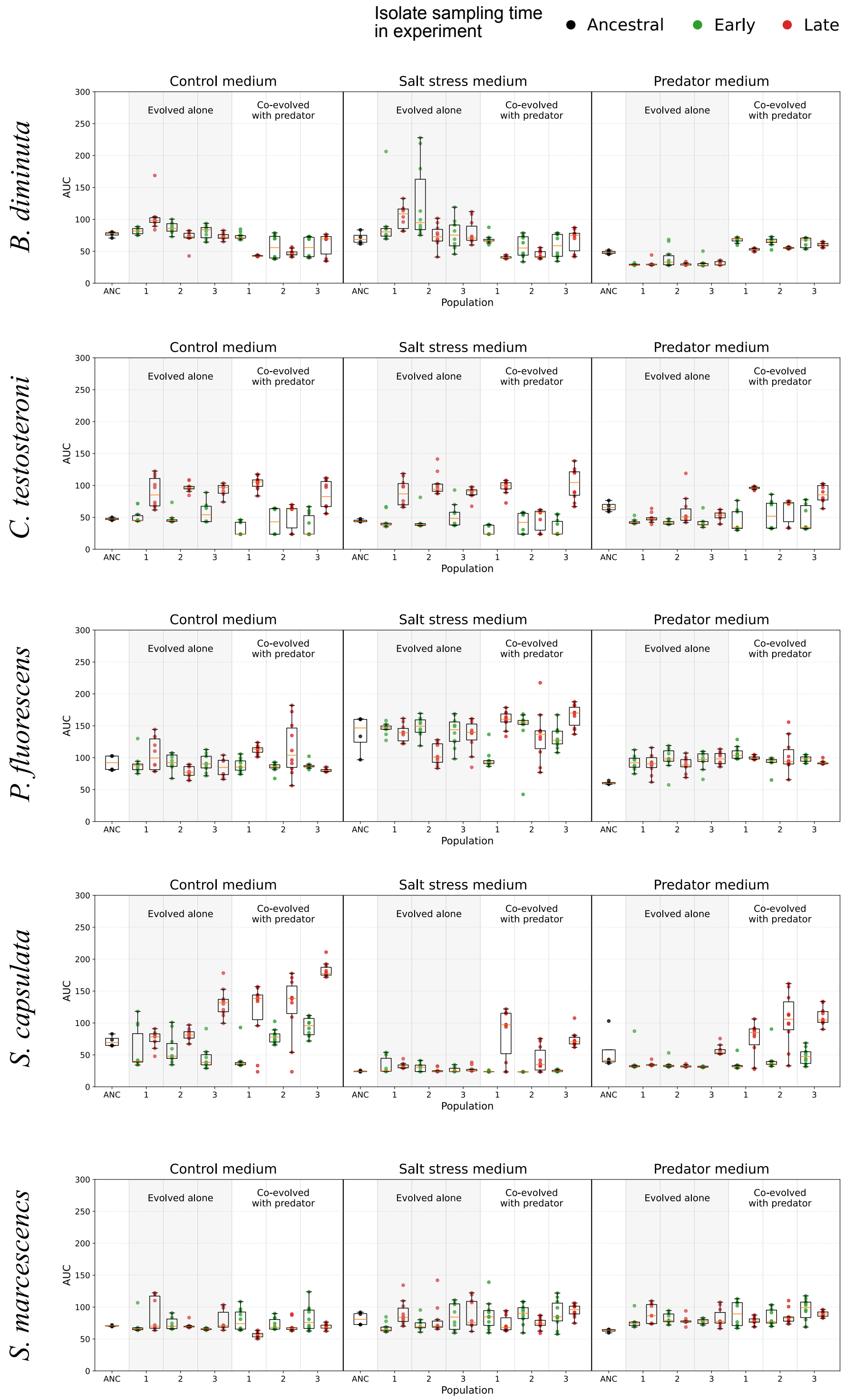


Figure 1: Area under the curve (AUC) of isolate growth across experiments. Each row represents one species in our experiment and is divided into three sections — one of each growth environment (control, salt-stress and predator mediums). Each medium section is further divided into ancestral (”ANC”), evolved alone and coevolved with predator sections to refer to the coevolutionary history of isolates. In each replicate population (marked on the X-axis), isolates sampled at the two sampling timings are shown and colored separately (early – green, late – red). Ancestral isolates are colored black.

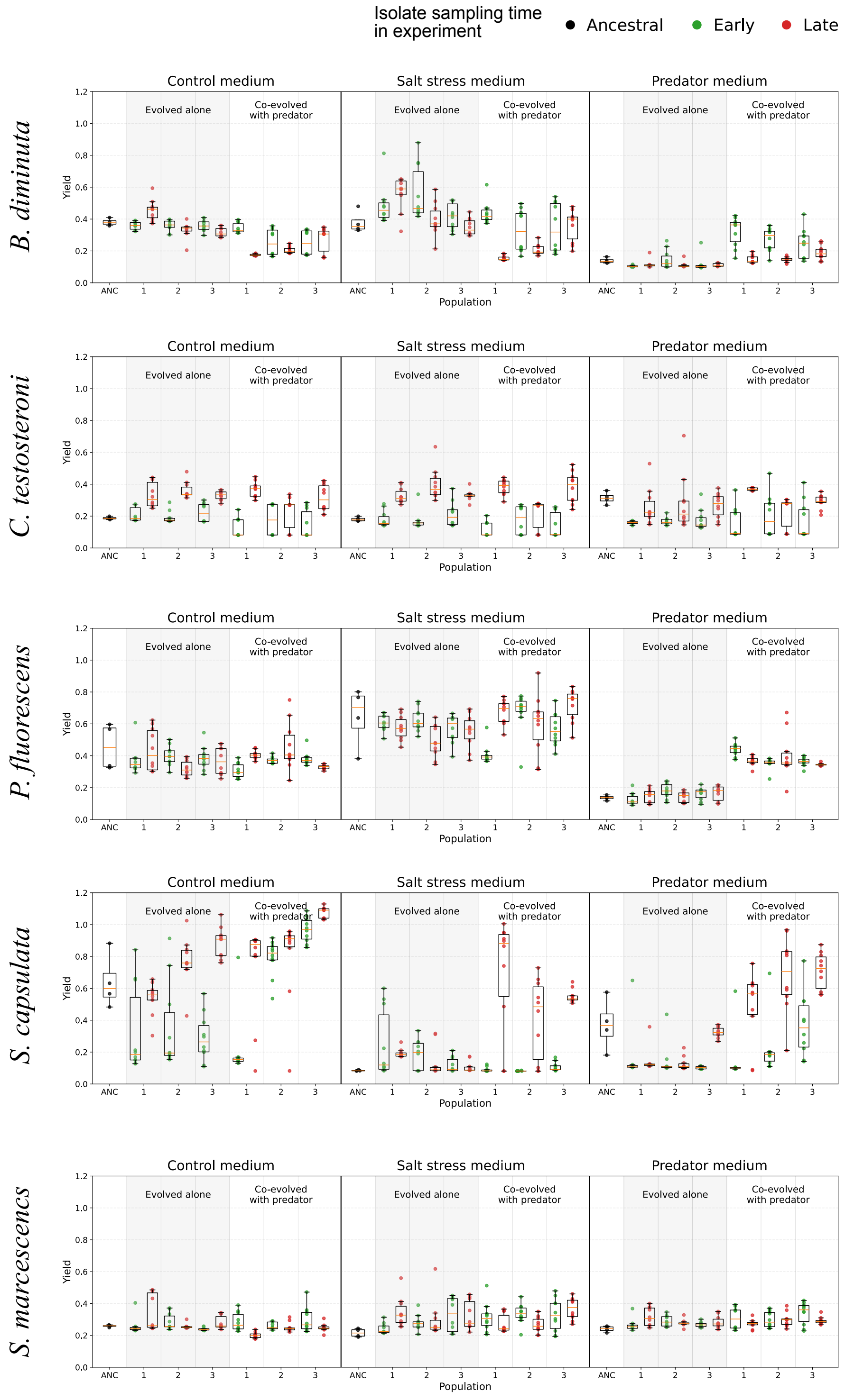


Figure 2: Yield of isolate growth across experiments. Each row represents one species in our experiment and is divided into three sections — one of each growth environment (control, salt-stress and predator mediums). Each medium section is further divided into ancestral ("ANC"), evolved alone and coevolved with predator sections to refer to the coevolutionary history of isolates. In each replicate population (marked on the X-axis), isolates sampled at the two sampling timings are shown and colored separately (early – green, late – red). Ancestral isolates are colored black.

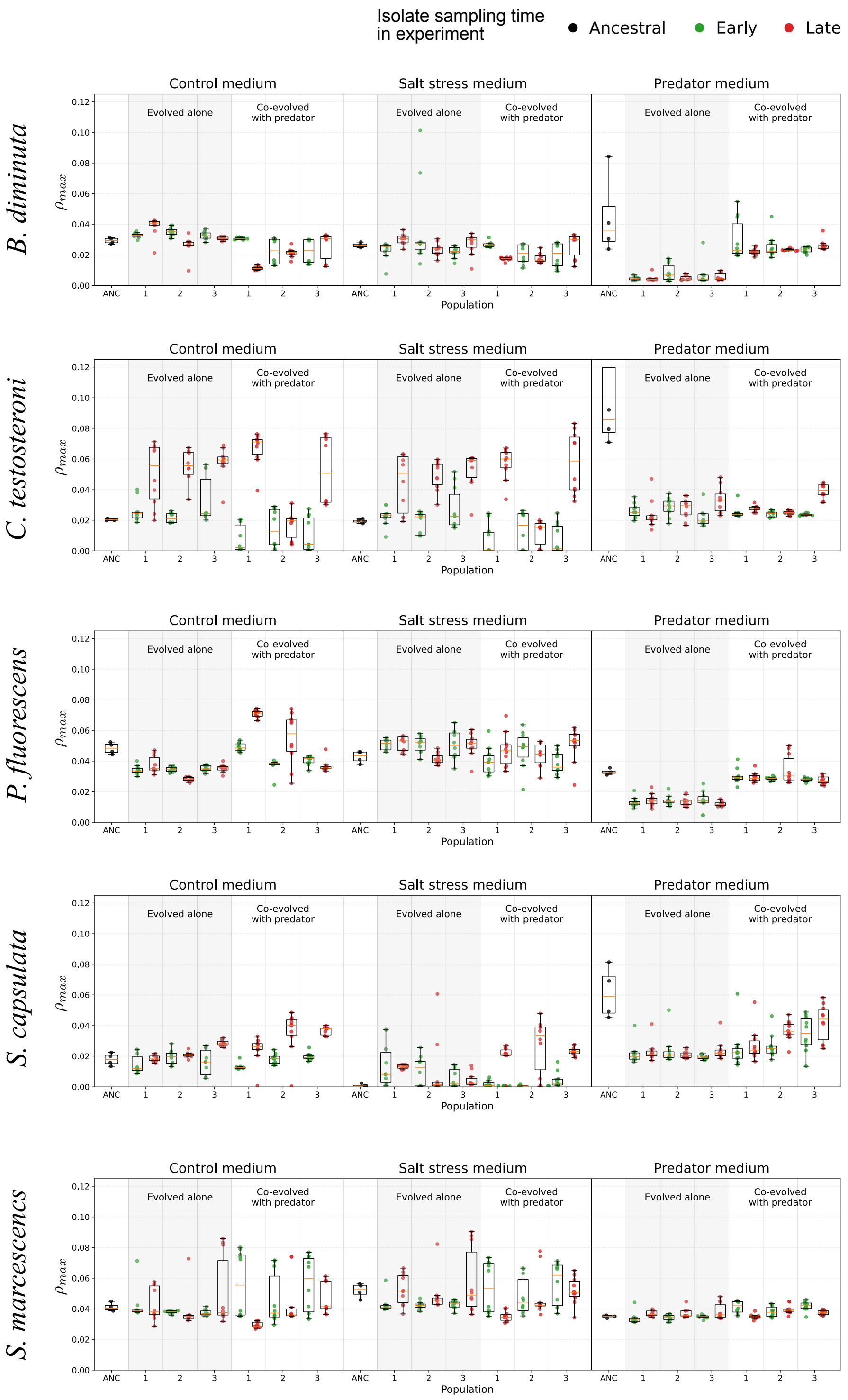


Figure 3: Maximum per-capita growth rate $\rho_{max}$ of isolate growth across experiments. Each row represents one species in our experiment and is divided into three sections — one of each growth environment (control, salt-stress and predator mediums). Each medium section is further divided into ancestral ("ANC"), evolved alone and coevolved with predator sections to refer to the coevolutionary history of isolates. In each replicate population (marked on the X-axis), isolates sampled at the two sampling timings are shown and colored separately (early – green, late – red). Ancestral isolates are colored black.

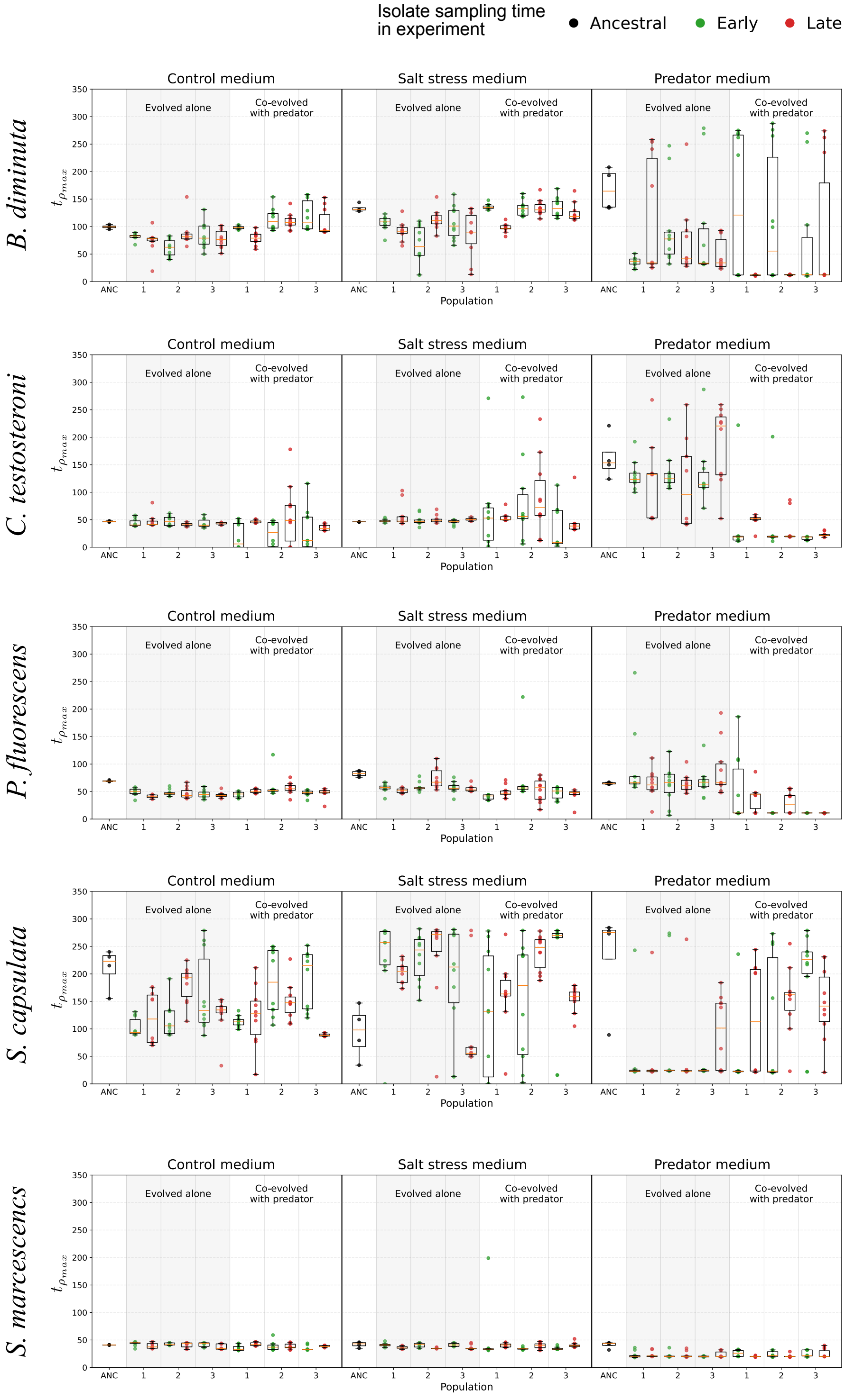


Figure 4: Time to the maximum per-capita growth rate $t_{\rho_{max}}$ of isolate growth across experiments. Each row represents one species in our experiment and is divided into three sections — one of each growth environment (control, salt-stress and predator mediums). Each medium section is further divided into ancestral ("ANC"), evolved alone and coevolved with predator sections to refer to the coevolutionary history of isolates. In each replicate population (marked on the X-axis), isolates sampled at the two sampling timings are shown and colored separately (early – green, late – red). Ancestral isolates are colored black.

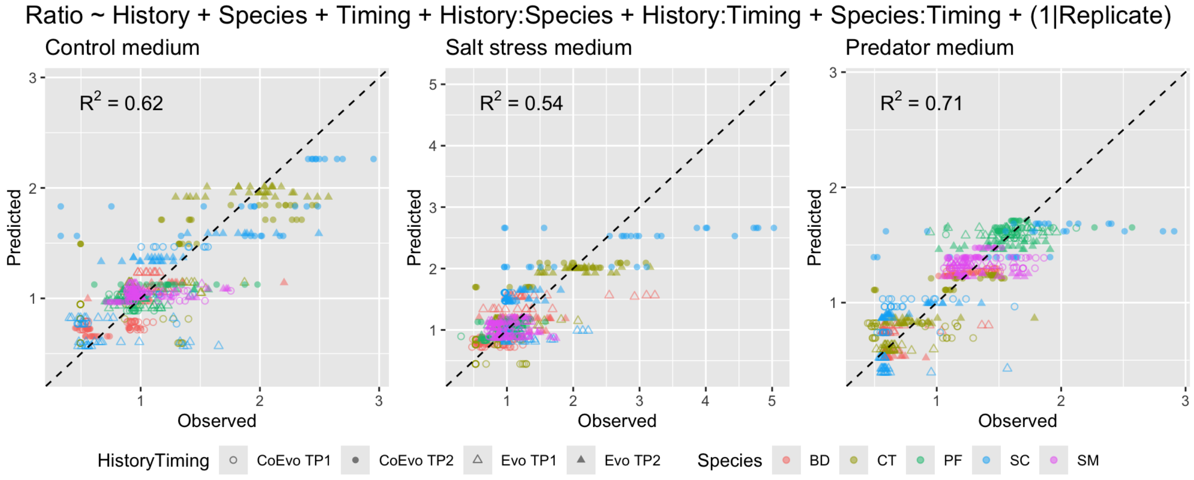


Figure 5: Scatter plot of observed vs. linear mixed model with three interactions between main terms fitted relative fitness values. Each sub-panel represents one of the three growth mediums (control, salt stress and predator). Model specification is given at the top ("Ratio": relative fitness, "History": evolutionary history, "Species": species identity, "Timing": isolate sampling timing, "Replicate": replicate population identifier, specific to a species within either evolutionary history). Evolutionary histories are marked as circles (coevolved prey) and triangles (evolved-alone prey), and these markers are filled depending on isolate sampling timing (empty: early sampling, filled: late sampling). Colors denote each species ("BD": *B. diminuta*, "CT": *C. testosteroni*, "PF": *P. fluorescens*, "SC": *S. capsulata*, "SM": *S. marcescens*). $R^2$ in each sub-panel indicates proportion of variance explained by the linear mixed model (i.e., goodness-of-fit).

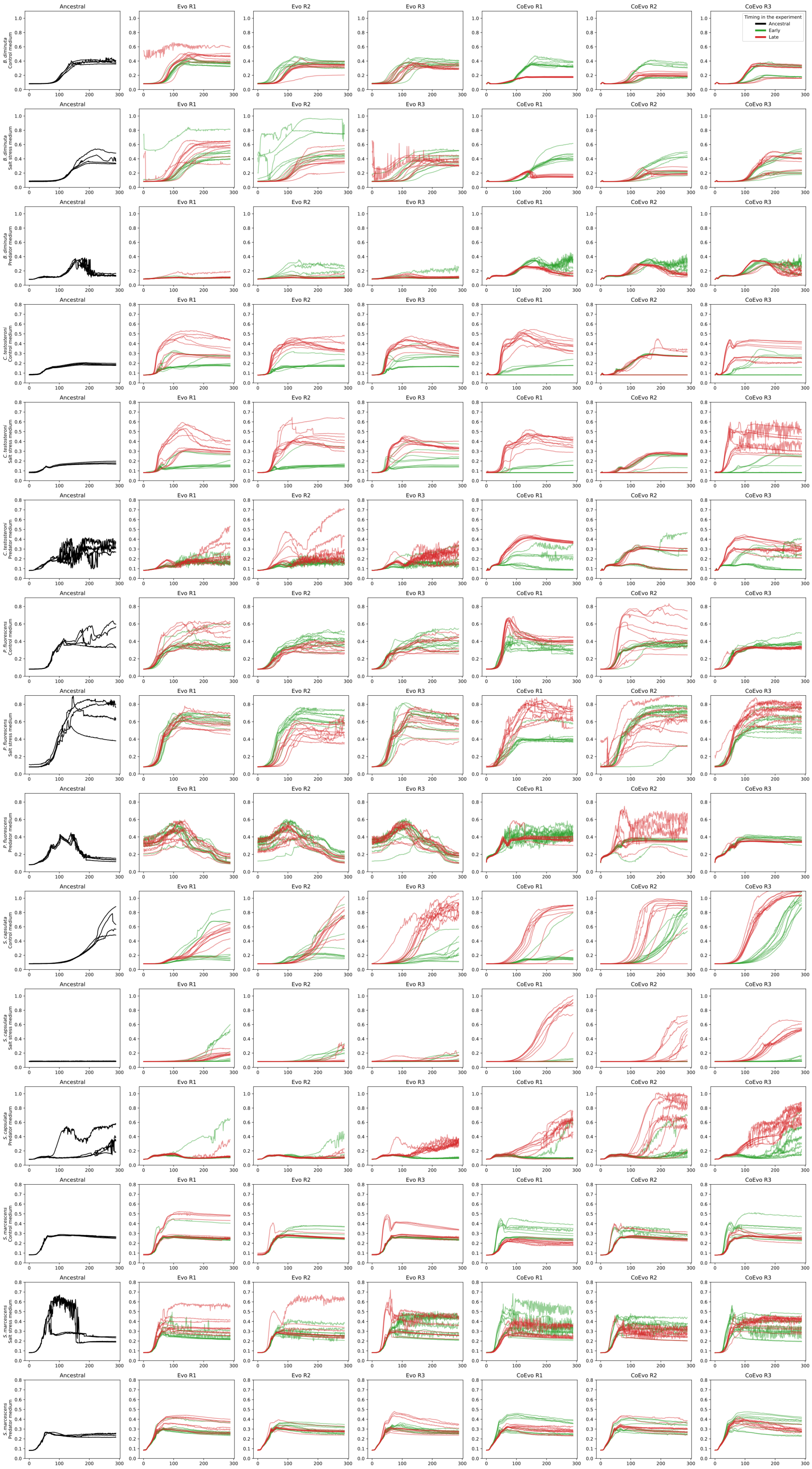


Figure 6: Phenotypic measurements (growth curves) of each study population. Each row represents one species in one growth medium (see y-axis label). First column represents ancestral isolate growth of the respective species, and subsequent columns are study populations ("Evo": evolved-alone; "CoEvo": coevolved with ciliate). Green colored curves indicate early sampling isolates, and red curves indicate late isolates.

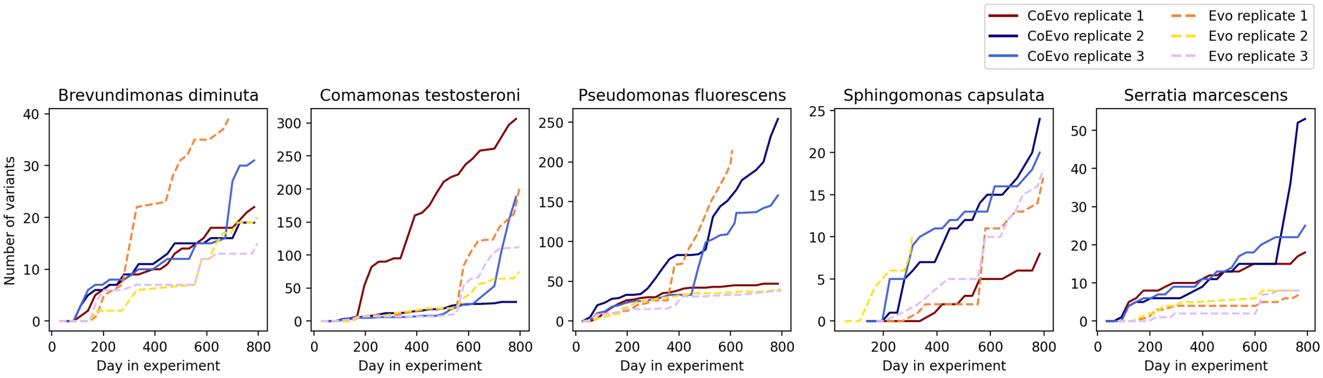


Figure 7: Mutation accumulation over the course of the experiment. "CoEvo" refers to prey populations that have coevolved with the predator over the course of the long-term experiment, and "Evo" denotes prey populations that were evolved alone.

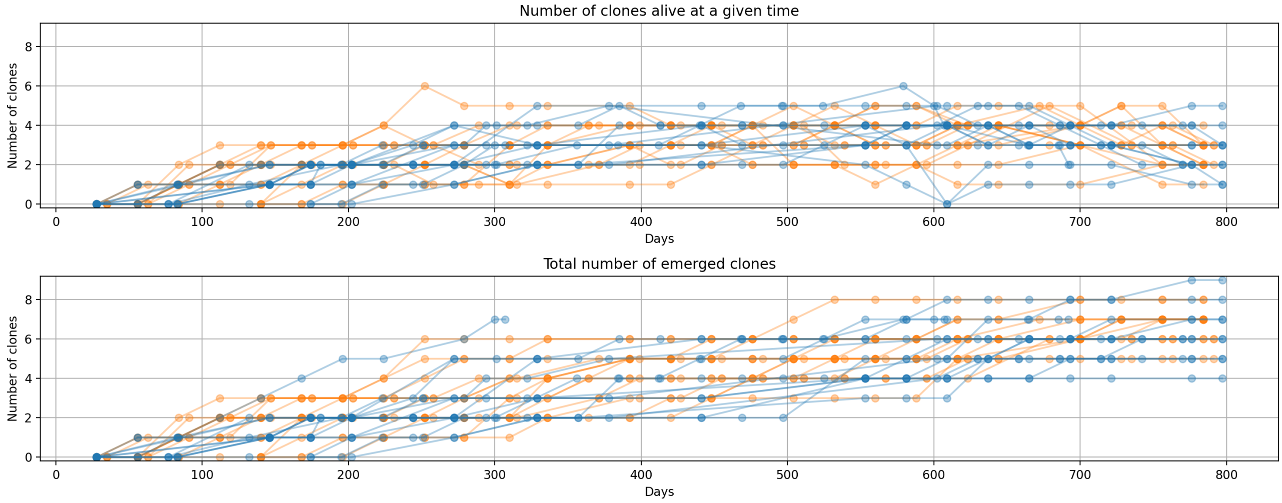


Figure 8: Changes in the number of clones over the course of the long-term experiment. Each line represents one population (orange lines – populations that coevolved with ciliate, blue lines – populations that evolved alone). **Top panel:** number of alive clones (i.e., clone frequency > 0.1) at any given time in the experiment. **Bottom panel:** total number of clones accumulated over the course of the experiment.

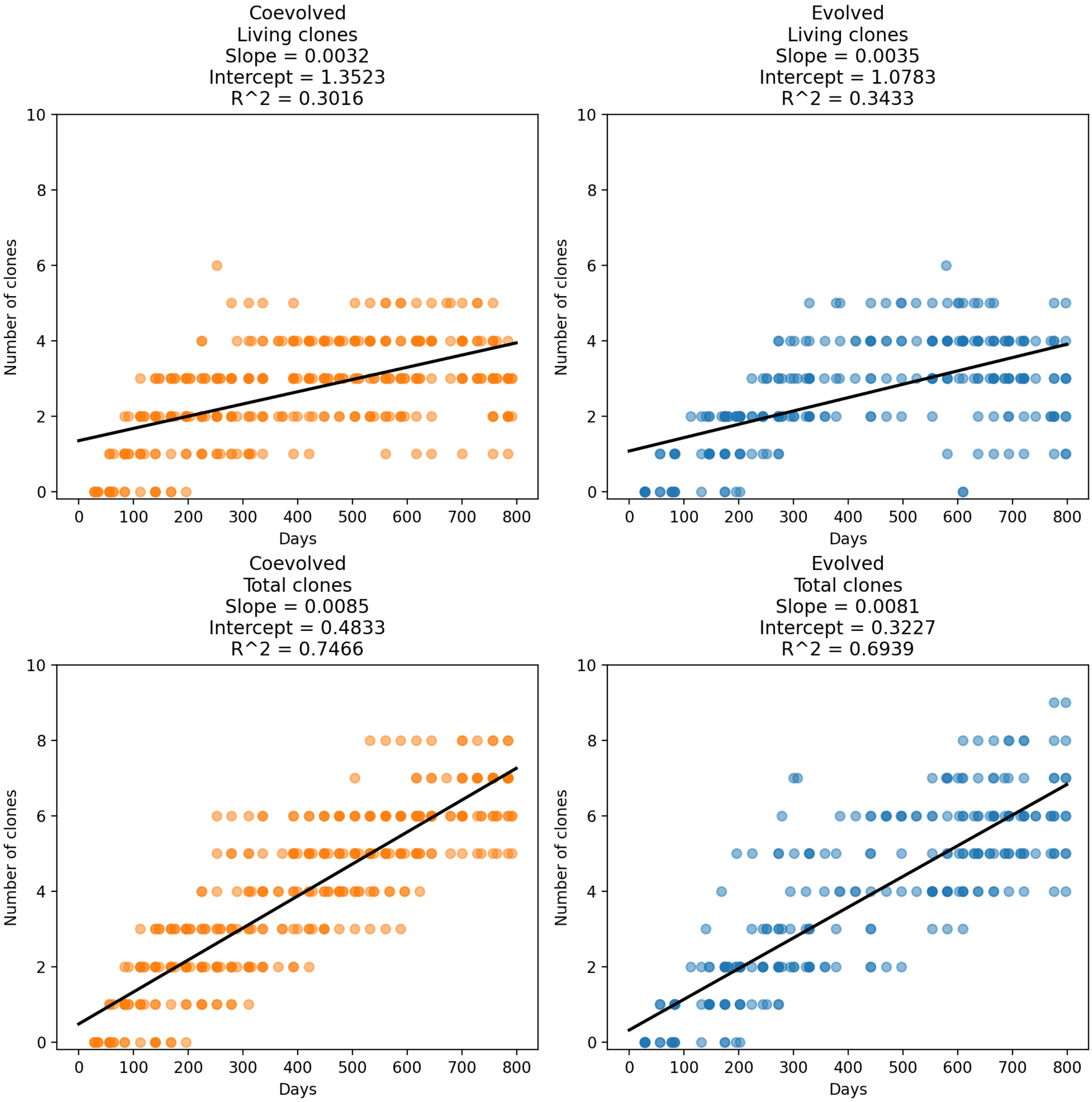


Figure 9: Tests results of linear regression on living and total accumulated clone counts (see Supplementary Fig. 8). Orange color denotes populations that coevolved with ciliate, blue color – populations that evolved alone. Subfigure titles include slope, intercept and the coefficient of determination $R^2$.

#### *B. diminuta* (coevolved with predator)

**Population 1** **Population 2** **Population 3**

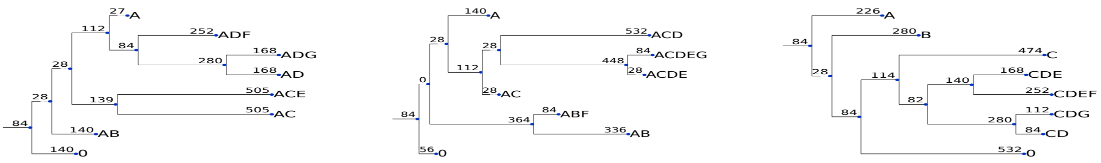


Number of days: 784 Number of days: 784 Number of days: 784

#### *C. testosteroni* (coevolved with predator)

**Population 1** **Population 2** **Population 3**

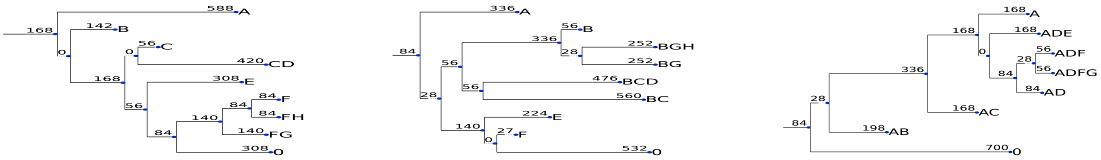


Number of days: 784 Number of days: 784 Number of days: 784

#### *P. fluorescens* (coevolved with predator)

**Population 1** **Population 2** **Population 3**

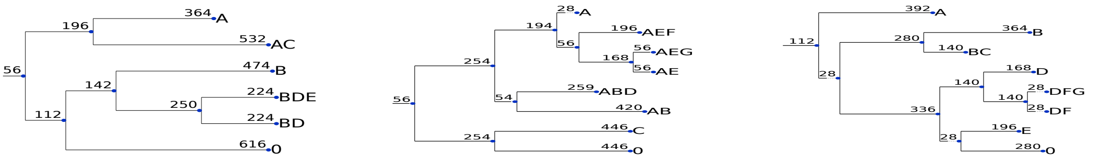


Number of days: 784 Number of days: 784 Number of days: 784

#### *S. capsulata* (coevolved with predator)

**Population 1** **Population 2** **Population 3**

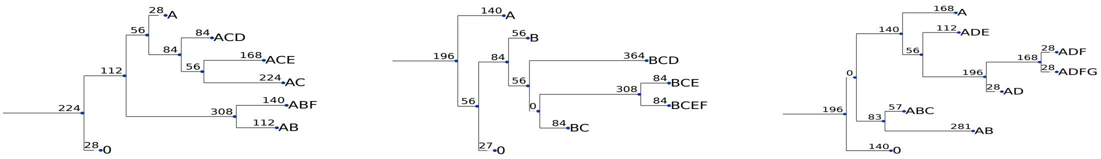


Number of days: 784 Number of days: 784 Number of days: 784

#### *S. marcescens* (coevolved with predator)

**Population 1** **Population 2** **Population 3**

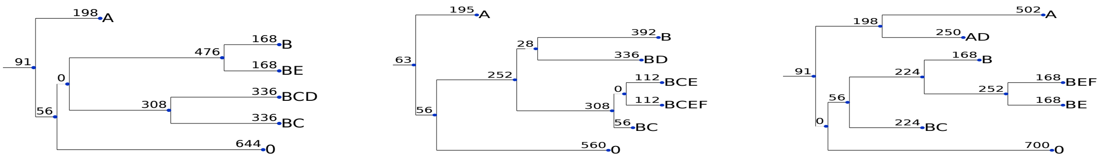


Number of days: 791 Number of days: 791 Number of days: 791

Figure 10: Phylogenetic trees for study populations that were coevolving with the ciliate.

#### *B. diminuta* (evolved alone)

**Population 1** **Population 2** **Population 3**

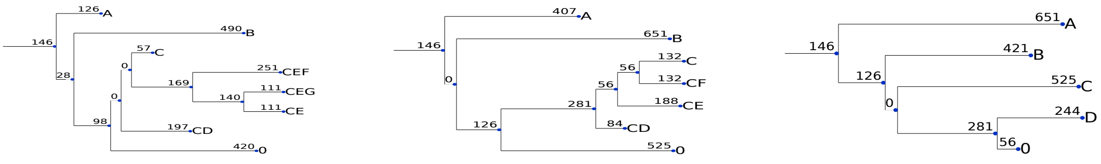

Number of days: 692 Number of days: 797 Number of days: 797

#### *C. testosteroni* (evolved alone)

**Population 1** **Population 2** **Population 3**

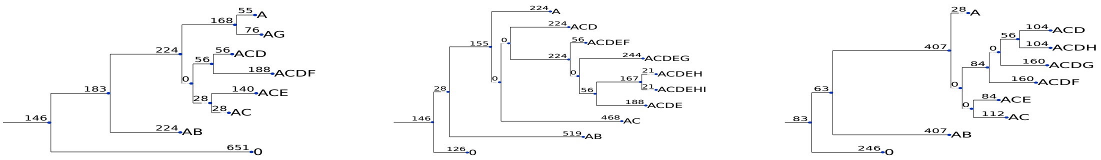

Number of days: 797 Number of days: 797 Number of days: 797

#### *P. fluorescens* (evolved alone)

**Population 1** **Population 2** **Population 3**

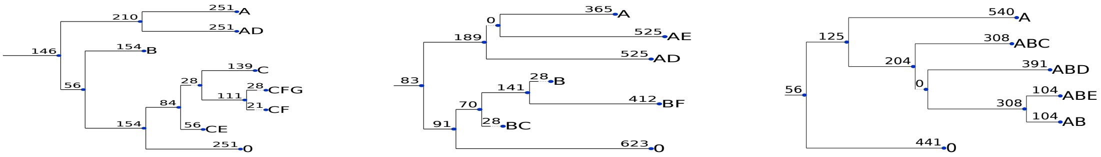

Number of days: 607 Number of days: 797 Number of days: 797

#### *S. capsulata* (evolved alone)

**Population 1** **Population 2** **Population 3**

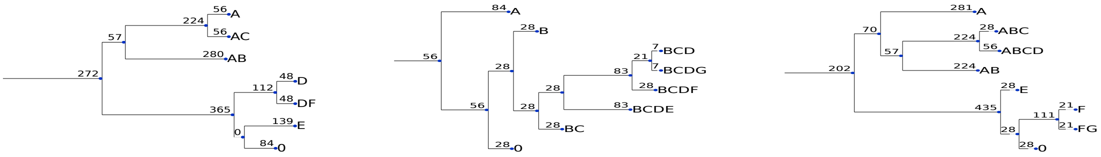

Number of days: 797 Number of days: 307 Number of days: 797

#### *S. marcescens* (evolved alone)

**Population 1** **Population 2** **Population 3**

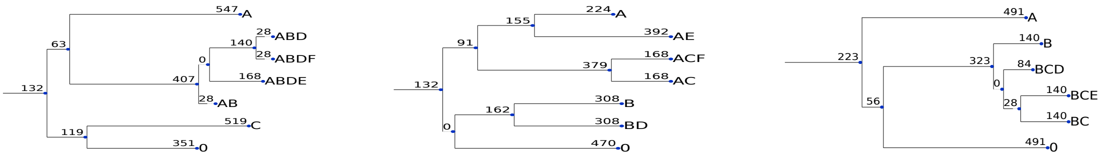

Number of days: 770 Number of days: 770 Number of days: 770

Figure 11: Phylogenetic trees for study populations that were evolving alone.

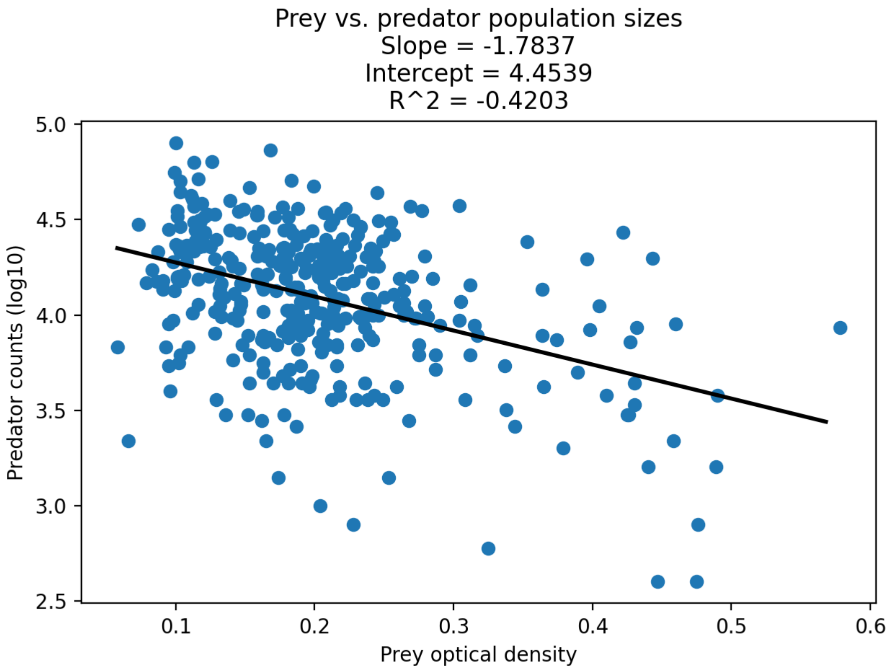


Figure 12: Scatter-plot of prey optical density versus predator counts (log10) and linear regression results. Slope, intercept and the coefficient of determination $R^2$ are reported in the figure title.

### Supplementary Tables

| | Evolved alone | | | | Coevolved | | | |
|---|---|---|---|---|---|---|---|---|
| **Species** | **Pop. 1** | **Pop. 2** | **Pop. 3** | **Mean** | **Pop. 1** | **Pop. 2** | **Pop. 3** | **Mean** |
| *B. diminuta* | 39 | 20 | 15 | 24.7 | 22 | 19 | 31 | 24 |
| *C. testosteroni* | 201 | 75 | 112 | 129.3 | 306 | 29 | 188 | 174.3 |
| *P. fluorescens* | 215 | 40 | 38 | 97.7 | 47 | 254 | 158 | 153 |
| *S. capsulata* | 17 | 10 | 18 | 15 | 8 | 24 | 20 | 17.3 |
| *S. marcescens* | 7 | 8 | 8 | 7.7 | 18 | 53 | 25 | 32 |

Table 1: Accumulated variant counts per studied population.

### Statistical testing of phylogenetic trees

| **Metric** | **p value** |
|---|---|
| Sum | 0.053 |
| Mean | 0.431 |
| Variance | 0.901 |
| Number of leaves | 0.528 |
| Maximum depth | 0.759 |
| Average depth | 0.754 |
| Sackin index | 0.752 |
| Colless index | 0.928 |
| B1 index | 0.285 |
| ${}^0D_N$ | 0.201 |
| ${}^1D_N$ | 0.609 |
| ${}^0D_S$ | 0.134 |
| ${}^1D_S$ | 0.059 |
| ${}^0D_L$ | 0.645 |
| ${}^1D_L$ | 0.48 |
| ${}^1J_N$ | 0.179 |
| ${}^1J_S$ | 0.168 |
| ${}^1J_L$ | 0.206 |

Table 2: Resulting uncorrected p-value for each phylogenetic tree measurement using ANCOVA (pooled data with species indicator as a covariate). Sum, mean, variance are computed directly from tree branch lengths. ${}^0D_N$, ${}^1D_N$, ${}^0D_S$, ${}^1D_S$, ${}^0D_L$, ${}^1D_L$, ${}^1J_N$, ${}^1J_S$, ${}^1J_L$ are the RUI tree indices (Noble and Verity 2023)

| Species \ Metric | Sum | Mean | Variance | Clusters | Max depth | Avg depth | Sackin ind. | Colless ind. | B1 ind. |
|---|---|---|---|---|---|---|---|---|---|
| *B. diminuta* | 0.594 | 0.348 | 0.3 | 0.205 | 0.725 | 0.487 | 0.321 | 0.628 | 0.004* |
| *C. testosteroni* | 0.306 | 0.114 | 0.096 | 0.643 | 0.643 | 0.448 | 0.458 | 0.485 | 0.921 |
| *P. fluorescens* | 0.336 | 0.696 | 0.727 | 0.725 | 0.643 | 0.807 | 0.898 | 0.693 | 0.259 |
| *S. capsulata* | 0.591 | 0.532 | 0.812 | 0.519 | 0.468 | 0.661 | 0.948 | 0.575 | 0.433 |
| *S. marcescens* | 0.494 | 0.413 | 0.492 | 1.0 | 0.678 | 0.398 | 0.547 | 0.51 | 0.926 |

Table 3: Resulting uncorrected p-value for each phylogenetic tree measurement using ANOVA (each species tested separately). Sum, mean, variance are computed directly from tree branch lengths. An asterisk next to a p-value marks the significance of the results given 95% confidence level.

| Species \ Metric | $^0D_N$ | $^1D_N$ | $^1J_N$ | $^0D_S$ | $^1D_S$ | $^1J_S$ | $^0D_L$ | $^1D_L$ | $^1J_L$ |
|---|---|---|---|---|---|---|---|---|---|
| *B. diminuta* | 0.054 | 0.008* | 0.644 | 0.298 | 0.189 | 0.379 | 0.595 | 0.416 | 0.639 |
| *C. testosteroni* | 0.498 | 0.344 | 0.194 | 0.793 | 0.47 | 0.292 | 0.288 | 0.245 | 0.182 |
| *P. fluorescens* | 0.777 | 0.806 | 0.912 | 0.574 | 0.557 | 0.661 | 0.577 | 0.823 | 0.938 |
| *S. capsulata* | 0.186 | 0.338 | 0.622 | 0.47 | 0.369 | 0.586 | 0.554 | 0.741 | 0.658 |
| *S. marcescens* | 0.137 | 0.586 | 0.891 | 0.786 | 0.787 | 0.499 | 0.767 | 0.931 | 0.867 |

Table 4: Resulting uncorrected p-value for each of the phylogenetic tree RUI indices (Noble and Verity 2023) using ANOVA (each species tested separately). An asterisk next to a p-value marks the significance of the results given 95% confidence level.

### Recurrently targeted gene statistics

| Gene \ Tested between | **Experiment** | ***B. diminuta*** | ***C. testosteroni*** | ***P. fluorescens*** | ***S. capsulata*** | ***S. marcescens*** |
|---|---|---|---|---|---|---|
| *barA* | 0.021983* | 0.4254 | 0.4254 | 0.4254 | 0.4254 | 0.000021* |
| *cheB* | 0.021983* | 0.235688 | 0.235688 | 0.000059* | 0.235688 | 0.235688 |
| *dppB* | 0.021983* | 0.235688 | 0.235688 | 0.000059* | 0.235688 | 0.235688 |
| *flhC* | 0.021983* | 0.235688 | 0.235688 | 0.235688 | 0.235688 | 0.000059* |
| *lptA* | 0.031923* | 0.269164 | 0.779762 | 0.102679 | 0.269164 | 0.779762 |
| *puuR* | 0.021983* | 0.4254 | 0.4254 | 0.000021* | 0.4254 | 0.4254 |
| *rcsC* | 0.028205* | 0.446983 | 0.012335* | 0.135119 | 0.065135 | 0.446983 |
| *relA* | 0.021983* | 0.4254 | 0.000021* | 0.4254 | 0.4254 | 0.4254 |
| *rpoB* | 0.011444* | 0.082328 | 0.082328 | 0.373633 | 0.654406 | 0.373633 |
| *rpoC* | 0.016356* | 0.212047 | 1.0 | 1.0 | 1.0 | 0.212047 |
| *sasA* | 0.018005* | 0.835325 | 0.40887 | 0.40887 | 0.070708 | 0.029696* |

Table 5: Genes that are statistically significantly recurring between experiments (with ciliate and without ciliate) across all species ("Experiment“ column) and within a given species (columns with species names), as per ANCOVA testing. Asterisks mark significant (uncorrected) p-values at a 95% confidence level.
